# Is faster-X adaptation due to large-effect mutations? An empirical test of a new theory

**DOI:** 10.64898/2026.02.06.704462

**Authors:** Filip Ruzicka, Beatriz Vicoso

## Abstract

A widespread observation in molecular evolution is that X-linked genes evolve faster—and often adapt faster—than autosomal genes. Three main ideas have been put forward to explain “faster-X” adaptation. The first is that new beneficial mutations are typically partially recessive with respect to fitness, leading to more efficient selection in hemizygous males. The second is that sex differences in recombination, or sex differences in variance in reproductive success, enhance the effective population size on the X chromosome relative to autosomes. Both ideas have limitations: the first conflicts with theories of dominance, in which beneficial mutations are proposed to be partially dominant, while the second only applies to some taxa. Here we tested a third, newly-proposed theory, in which mutations with large “scaled phenotypic effects” (i.e., large effects relative to the distance to a phenotypic optimum) experience more positive selection on the X. Specifically, we used three proxies for scaled phenotypic effects—amino-acid dissimilarity, sequence conservation, and gene age—and estimated X and autosomal rates of adaptation of nonsynonymous mutations across effect-size classes. We did this in three lineages with a well-documented faster-X: *Drosophila melanogaster*, *Mus musculus*, and *Homo sapiens*. As expected, we found that large-effect mutations were more likely to be under purifying selection and less likely to be under positive selection than small-effect mutations. However, contrary to the new theory, faster-X adaptation was not enriched among large-effect mutations. Overall, our results highlight an unresolved gap between patterns of sex chromosome evolution and theories of dominance, and the need for more direct empirical data on the dominance of new beneficial mutations.

**Significance statement:** In many species, genes on the X chromosome undergo more adaptation than autosomal genes, but it is not clear why this pattern of “faster-X” occurs. One possibility is that recessive beneficial mutations drive the effect, but theories of dominance propose instead that beneficial mutations should be dominant. Meanwhile, alternative explanations (e.g., sex differences in recombination or sex differences in variance in reproductive success) can only apply to some taxa. Here we tested a new idea, which proposes that the effect is driven by mutations with large effects. However, data from fruit flies, humans and mice did not provide support for the new theory. Our work highlights an unresolved gap between patterns of sex chromosome evolution and theories of dominance.

## Introduction

Empirical tests of evolutionary theories require exceptional study systems in which model parameters are both measurable and allowed to vary. In species with genetic sex determination, X-linked and autosomal genes exhibit different modes of inheritance, generating fortuitous natural variation in many genetic parameters and permitting a wide range of inferences about evolution (Vicoso and Charlesworth 2006; Ellegren 2009; Wilson Sayres 2018). For example, X-autosome contrasts have been used to estimate sex differences in mutation rates (Miyata et al. 1987; Wilson Sayres and Makova 2011; Connallon et al. 2022), breeding ratios (Hammer et al. 2010; Lohmueller et al. 2010) and migration rates (Juric et al. 2016; Goldberg et al. 2017; Osada et al. 2020; Chevy et al. 2023). X-autosome contrasts have also been used to infer the dominance coefficients of deleterious mutations (Avery 1984; Mallet et al. 2011; Veeramah et al. 2014; Ruzicka et al. 2021), beneficial mutations (Haldane 1927; Charlesworth et al. 1987; Orr 2010; Veeramah et al. 2014; Charlesworth et al. 2018) and sexually antagonistic mutations (Rice 1984; Fry 2010; Frank and Patten 2020; Ruzicka and Connallon 2020; Ruzicka and Connallon 2022). And they are key components of hypotheses about genome evolution (Meisel et al. 2009; Vibranovski et al. 2009; Toups and Hahn 2010), lifespan (Brengdahl et al. 2018; Sultanova et al. 2018; Connallon et al. 2022; Sultanova et al. 2023: 202) and speciation (Turelli and Orr 1995; Turelli and Orr 2000; Masly and Presgraves 2007; Ellegren et al. 2012; Sankararaman et al. 2016; Presgraves 2018).

In the last 20 years, a consistent pattern has emerged from X-autosome comparisons (Meisel and Connallon 2013; Charlesworth et al. 2018; McDonough et al. 2024): namely, that functional sequence divergence is elevated among X-linked genes, relative to autosomal genes (Baines et al. 2008; Vicoso et al. 2013; Sackton et al. 2014; Wright et al. 2015; Jaquiéry et al. 2018; Bechsgaard et al. 2019; Mongue et al. 2022; Darolti et al. 2023; Mongue and Baird 2024). This pattern of “faster-X (or faster-Z) evolution” can either be driven by less effective purifying selection (stronger drift), or more effective positive selection, on the X chromosome relative to autosomes (Vicoso and Charlesworth 2009; Mank et al. 2010). In some cases, the contributions of positive selection to faster-X evolution have been isolated, using McDonald-Kreitman (MK) tests that compare polymorphism and divergence (McDonald and Kreitman 1991; Eyre-Walker and Keightley 2009; Galtier 2016), or population genomic signals of selective sweeps. A pattern of “faster-X (or faster-Z) adaptation” often emerges (see (McDonough et al. 2024) for a recent review), as in the case of *Heliconius* butterflies (Pinharanda et al. 2019), *Drosophila* fruit flies (Ávila et al. 2014; Campos et al. 2014; Garrigan et al. 2014; Harris et al. 2024), apes (Hvilsom et al. 2012; Veeramah et al. 2014; Dutheil et al. 2015) and rodents (Carneiro et al. 2012; Kousathanas et al. 2014). Some species, often ZW systems, show no significant X/Z-autosome differences (Rousselle et al. 2016; Hayes et al. 2020; Baird et al. 2025; Chase et al. 2025), though the only clear example of faster-autosome adaptation to date is the blood fluke *Schistosoma japonicum*, in which Z-linked effective population sizes (*N_e_*) are drastically reduced (Mrnjavac and Vicoso 2025). Moreover, an analogous pattern of “faster-haploid” adaptation, relative to diploids, has been repeatedly reported in experimental populations of the budding yeast *Saccharomyces cerevisiae* (Zeyl et al. 2003; Gerstein et al. 2011; Marad et al. 2018).

Three main ideas have been put forward to explain faster-X *adaptation*. The first and most commonly invoked idea is that beneficial mutations are on average partially recessive with respect to fitness (Charlesworth et al. 1987; Orr and Otto 1994). Partially recessive beneficial mutations that arise on the X are immediately exposed to positive selection in hemizygous (haploid) males, which—assuming adaptation proceeds from new mutations, equivalent fitness effects of homozygotes and hemizygotes, and *N_eX_* = ¾ *N_eA_*—enhances their substitution rates relative to equivalent autosomal mutations (Avery 1984; Charlesworth et al. 1987). This faster-X prediction when beneficial mutations are recessive is further exacerbated when selection is stronger in males than in females (Charlesworth et al. 1987; Meisel and Connallon 2013). This hypothesis is appealing because the dominance of mutations might be relatively similar across taxa, thus providing a general explanation for faster-X adaptation. However, the hypothesis also conflicts with theories of dominance, which propose that new beneficial mutations are on average partially dominant and/or overdominant for fitness (Kacser and Burns 1981; Manna et al. 2011; Sellis et al. 2011; McDonough and Connallon 2023; Ruzicka et al. 2025). If theories of dominance are correct, faster-autosome adaptation is to be expected, since autosomes reap the benefits of elevated mutational input while suffering minimal reductions in fixation probability due to incomplete dominance (Charlesworth et al. 1987; Meisel and Connallon 2013). Theories of dominance are widely supported by the observation that spontaneous deleterious mutations, especially strongly deleterious ones (Charlesworth 1979; Orr 1991; Agrawal and Whitlock 2011), are partially recessive (Manna et al. 2011). Direct data on the dominance of beneficial mutations are much sparser, and often biased by differences in fixation probability of dominant vs. recessive mutations (i.e., “Haldane’s sieve”), resulting in highly variable estimates (Bourguet and Raymond 1998; Anderson et al. 2004; Gerstein et al. 2011; Ronfort and Glémin 2013; Gerstein et al. 2014; Sellis et al. 2016; Marad et al. 2018; Aggeli et al. 2022).

A second idea is that sex differences in variance in reproductive success, or sex differences in recombination rate, generate conditions where *N_eX_* > ¾ *N_eA_*. For example, high variances in reproductive success tend to reduce *N_e_* (Charlesworth 2009), and such reductions are weaker on the X when variance in reproductive success is higher in males than females (Charlesworth 2001; Vicoso and Charlesworth 2009), due to the X chromosome’s female-biased transmission. Similarly, Hill-Robertson interference between mutations can be modelled as a reduction in *N_e_* (Charlesworth 2009), and this reduction is weaker on the X in species where male recombination is less frequent than female recombination (Charlesworth et al. 2018), again because of the X chromosome’s female-biased transmission. Thus, sex differences in variance in reproductive success or recombination rate can elevate *N_eX_*/*N_eA_*, potentially favouring faster-X adaptation even when beneficial mutations are partially dominant (Vicoso and Charlesworth 2009; Charlesworth et al. 2018). For example, the total absence of recombination in males of *Drosophila melanogaster* elevates X-autosome ratios of neutral diversity above 1 in some populations of this species (Charlesworth et al. 2018), which has been shown to contribute to faster-X adaptation (Campos et al. 2013). However, faster-X is also observed in *D. melanogaster* among pairs of genes with similar “effective” (sex-adjusted) recombination rates on the X and autosomes (Campos et al. 2013; Campos et al. 2014; Charlesworth et al. 2018). Moreover, signals of faster-X adaptation are found in species where X-autosome ratios of neutral diversity are near or below ¾ (Leffler et al. 2012), such as humans (Veeramah et al. 2014) and house mice (Kousathanas et al. 2014), which requires a different explanation.

Recently, a third idea to explain faster-X adaptation been proposed (McDonough et al. 2024), in which mutations with large “scaled phenotypic effects” (i.e., large phenotypic effects, relative to the distance to a phenotypic optimum) generate more positive selection on the X. As in classical faster-X theory, McDonough et al. (2024)’s theory assumes that adaptation proceeds by the sequential fixation of new mutations. However, they explicitly model the fitness effects of mutations using Fisher’s geometric model, in which new mutations pleiotropically affect many traits under stabilising selection towards a single phenotypic optimum. The rationale for faster-X adaptation is then as follows: (1) mutations can have large scaled phenotypic effects (i.e., either because phenotypic effects are large in absolute terms, or the optimum is nearby; see left vs. right panels of Fig. 1A) (Orr 1998); (2) large-effect mutations will typically undergo purifying selection (grey dots in Fig. 1A), but can sometimes express pleiotropic costs on other traits (red and yellow dots in Fig. 1A), leading to overdominance for fitness (Sellis et al. 2011; McDonough and Connallon 2023; Ruzicka et al. 2025); (3) overdominant mutations are subject to balancing selection and should remain polymorphic on autosomes, whereas the same mutations on the X chromosome can experience positive selection and rapidly become fixed (yellow dots in Fig. 1A) (Pamilo 1979). In other words, the model proposes that X-linkage converts a subset of mutations from evolving under balancing selection on autosomes to evolving under positive selection on the X (Fig. 1A), thereby generating an excess of adaptive substitutions on the X. A simple prediction of the model is that large-effect mutations should exhibit relatively high X-autosome ratios of rates of adaptive substitution compared to small-effect mutations because large-effect mutations are more susceptible to this “conversion” of balancing selection to positive selection on the X (Fig. 1B, C). McDonough et al. (2024)’s idea is appealing because it is consistent with leading theories of dominance (Manna et al. 2011), though it may still fail to generate faster-X if (for example) scaled phenotypic effects are consistently too small in real populations, or if scaled phenotypic effects cannot be accurately inferred.

**Figure 1.**
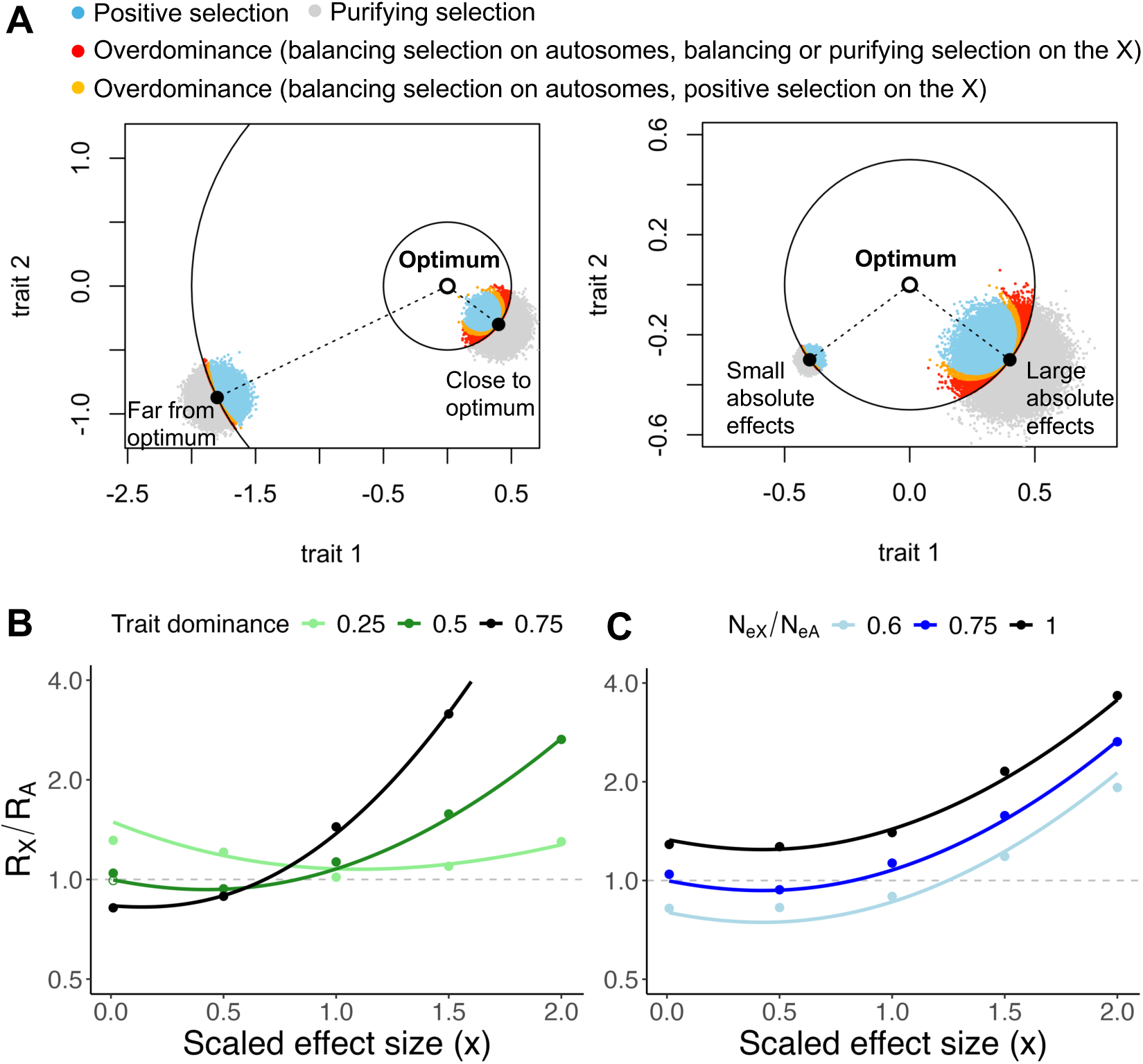
Theoretical background. **A.** Two-dimensional representation of adaptation in Fisher’s geometric model, showing an ancestral phenotype (black dots) and an optimum (white dot). Homozygous and heterozygous fitness effects are Gaussian functions of the distance to the optimum, and the points represent heterozygous fitness effects. The evolutionary dynamics of the mutations (positive selection, purifying selection, overdominance) depend on the mode of inheritance (autosomal or X-linked), expression in each sex, and trait dominance, with conditions outlined in McDonough et al. (2024). We show the case where mutations are expressed in both sexes, hemizygous and homozygous phenotypic effects are equivalent, and phenotypic effects of mutations are semi-dominant. The *scaled* phenotypic effect of a mutation (*x*) is a positive function of the absolute magnitude of the mutation (*r*) and trait dimensionality (√*n*), and a negative function of the distance to optimum (*z*). The left panel shows that mutations can have large *x* because they are close to the optimum (*z =* 0.5 left vs. *z = 2* right), or because they have large absolute magnitudes (*r* ≈ 0.15√*n* left vs. *r* ≈ 0.03√*n* right). This can generate overdominant balancing selection on autosomes (red and yellow points) but positive selection on the X (yellow points). **B.** Relative rates of adaptation on the X chromosome (*R_X_*) and autosomes (*R_A_*), on a log2 scale, plotted as a function of scaled phenotypic effect size (*x*) and phenotypic dominance. The curve is based on expressions for fixation probability in McDonough et al. (2024) (i.e., eq. (S21) for the X chromosome and eq. (S20) for autosomes), multiplied by 3/4 to account for enhanced mutational input on autosomes. Points represent Wright-Fisher simulations of positively selected mutations, with fitness effects generated in Fisher’s geometric model (*n* = 50, *z* = 1). Mutations were introduced sequentially with probability 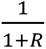 and 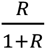 (for autosomes and the X, respectively, where 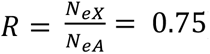 to account for differences in mutational input, and were run until at least 1000 fixations occurred on each chromosome type. We assumed *N_eA_* = 10^5^, 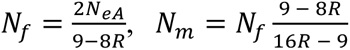, initial frequencies of 1/4*N*_*f*_ and 1/4*N*_*m*_ for autosomes, and 1/3*N*_*f*_and 1/3*N*_*m*_for the X (following the simulation procedure in Lasne et al. (2017)). Filled points denote simulations with fixed phenotypic dominance; the open point denotes a simulation where phenotypic dominance values were drawn from a uniform distribution between 0 and 1. **C.** Same as B, except that 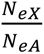 was varied, with phenotypic dominance fixed at 0.5.

Here, we tested McDonough et al. (2024)’s new hypothesis for faster-X adaptation by comparing X-autosome ratios of adaptive substitution rates, inferred from MK tests, among mutations of different effect sizes. Specifically, we used three proxies for the scaled phenotypic effects of mutations: differences in the physico-chemical properties of amino acid pairs (Grantham 1974; Bergman and Eyre-Walker 2019)—which are intended to capture variation in the absolute phenotypic effect of mutations (right panel of Fig. 1A)—and differences in sequence conservation (Vaser et al. 2016; Chen et al. 2022) and gene age (Domazet-Lošo et al. 2017; Moutinho et al. 2022)—which are intended to capture variation in the distance to the phenotypic optimum (left panel of Fig. 1A). We then examined whether signals of faster-X adaptation are stronger among large-effect than small-effect mutations. We applied this test to three model organisms covering a range of *N_eX_*/*N_eA_* ratios—above 1 in the fruit fly (*D. melanogaster*) (Ávila et al. 2014; Campos et al. 2014); below 0.75 in the house mouse (*Mus musculus castaneus*) (Kousathanas et al. 2014) and humans (*Homo sapiens*) (Veeramah et al. 2014)—and in which strong faster-X adaptation has previously been reported.

## Results

In *D. melanogaster*, one plausible explanation for faster-X adaptation is that genome-wide achiasmy in males elevates *N_eX_*/*N_eA_* to such a degree that there is no need to invoke recessive beneficial mutations (Vicoso and Charlesworth 2009; Charlesworth et al. 2018). We therefore began by examining the relationship between *N_eX_*/*N_eA_* ratios and the extent of faster-X adaptation in this species. Specifically, Hill-Robertson interference reduces *N_e_* in proportion to the extent of recombination, so that recombination rates can be used as proxies for local *N_e_* (Charlesworth 2009). We therefore estimated rates of adaptive substitution in 1Mb windows of recombination rate, after multiplying recombination rates by ½ on autosomes and 2/3 on the X to obtain “effective” recombination rates for each chromosome type and window. Rates of adaptive substitution for nonsynonymous sites were calculated as *ω*_*a*_, using a modern version of the McDonald-Kreitman test (see Materials and Methods), with *D. simulans* as the outgroup, polarising substitutions using *D. simulans* and *D. yakuba,* and correcting for possible biases due to segregating deleterious and beneficial mutations by fitting a DFE to unfolded site frequency spectra. As expected, effective recombination rates were higher on the X chromosome (mean cM/Mb = 1.97) than autosomes (mean cM/Mb = 1.09; see figure captions for results of all statistical tests). Importantly, however, a strong signal of faster-X adaptation was still observed in windows with the *same* effective recombination rate on the X and autosomes (Fig. 2). Thus, higher effective recombination rates on the X are not a sufficient explanation for faster-X adaptation in *D. melanogaster*.

**Figure 2.**
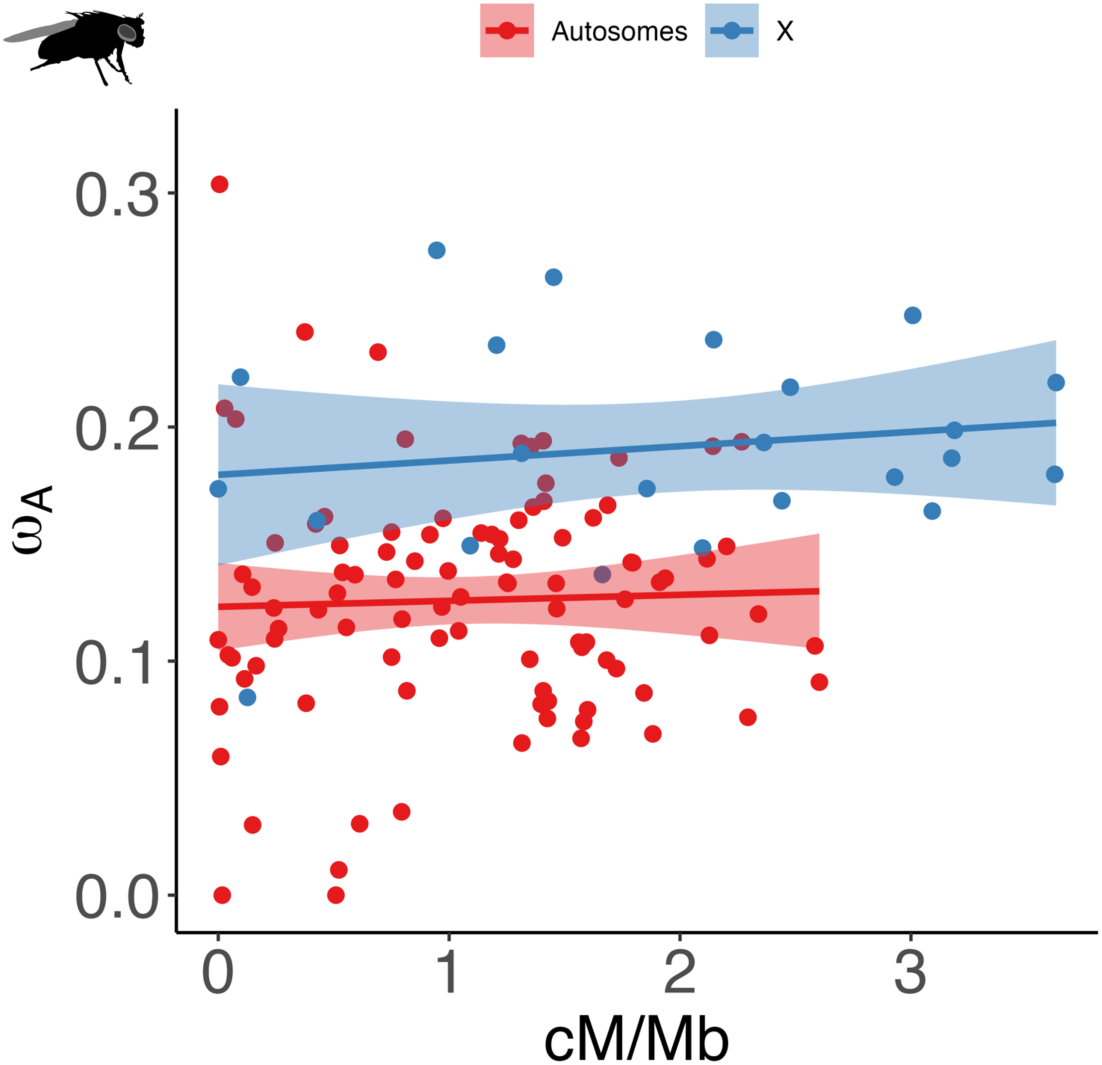
Effective recombination rates and relative rates of X-autosome adaptation in *D. melanogaster*. Estimates of *ω*_*a*_ for each 1Mb window of effective recombination rate (i.e., raw recombination rates in Comeron *et al*. (2012), multiplied by ½ on autosomes and 2/3 on the X chromosome, and averaged across 10 non-overlapping windows). Estimates of *ω*_*a*_ use the GammaExpo model to fit a DFE to segregating polymorphisms (see Materials and Methods). Negative estimates of *ω*_*a*_ were coded as zeroes. On average, recombination rate was higher on the X (Wilcoxon test, W = 72904, p<0.001). Residual *ω*_*a*_ (after correcting for recombination rate though linear regression; lines) was 0.034 on the X and –0.008 on autosomes (Wilcoxon test, W = 454, p<0.001).

To test the hypothesis that scaled phenotypic effects influence the extent of faster-X adaptation, we used three proxies for scaled phenotypic effects. First, we used differences in the physico-chemical properties of amino acid pairs, measured by their “Grantham’s distance” (Grantham 1974)—the rationale being that mutations with larger Grantham’s distances should have larger absolute phenotypic effects (see right panel of Fig. 1A) and thus larger scaled phenotypic effects. Second, we used differences in sequence conservation among nonsynonymous sites, measured by their “SIFT score” (Vaser et al. 2016), and differences in the age of genes that nonsynonymous sites fall in, with gene age inferred from phylogenetic patterns of gene orthology (Domazet-Lošo et al. 2017). The rationale here is that mutations in more conserved sites, or mutations in older genes, should be closer to their optimum, tending to increase their scaled phenotypic effects (see left panel of Fig. 1A).

To support the validity of our proxies, we took two approaches. We first correlated each metric with *π*_*N*_/*π*_*S*_, a measure of the efficiency of purifying selection (Castellano et al. 2018). In Fisher’s geometric model, mutations with larger scaled phenotypic effects (Fisher 1930; Orr 1998) are more likely to evolve under purifying selection, so that we expect large-effect mutations (inferred based on our proxies) to be under stronger purifying selection. Accordingly, we found that *π*_*N*_/*π*_*S*_ was strongly reduced among mutations with high amino-acid dissimilarity (Fig. 3A), in highly conserved sites (Fig. 3B), and in older genes (Fig. 3C) (see also (Bergman and Eyre-Walker 2019; Chen et al. 2022; Moutinho et al. 2022)). In the case of Grantham’s distances and gene age, this validation approach is robust, since the data used to generate both metrics and 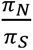 are independent. In the case of SIFT scores, they are not strictly independent from *π*_*N*_/*π*_*S*_, given that an evolutionary genetic model of purifying selection was used to generate these scores in the first place (Huber et al. 2020). This non-independence may therefore contribute to the correlation between SIFT categories and *π*_*N*_/*π*_*S*_.

**Figure. 3.**
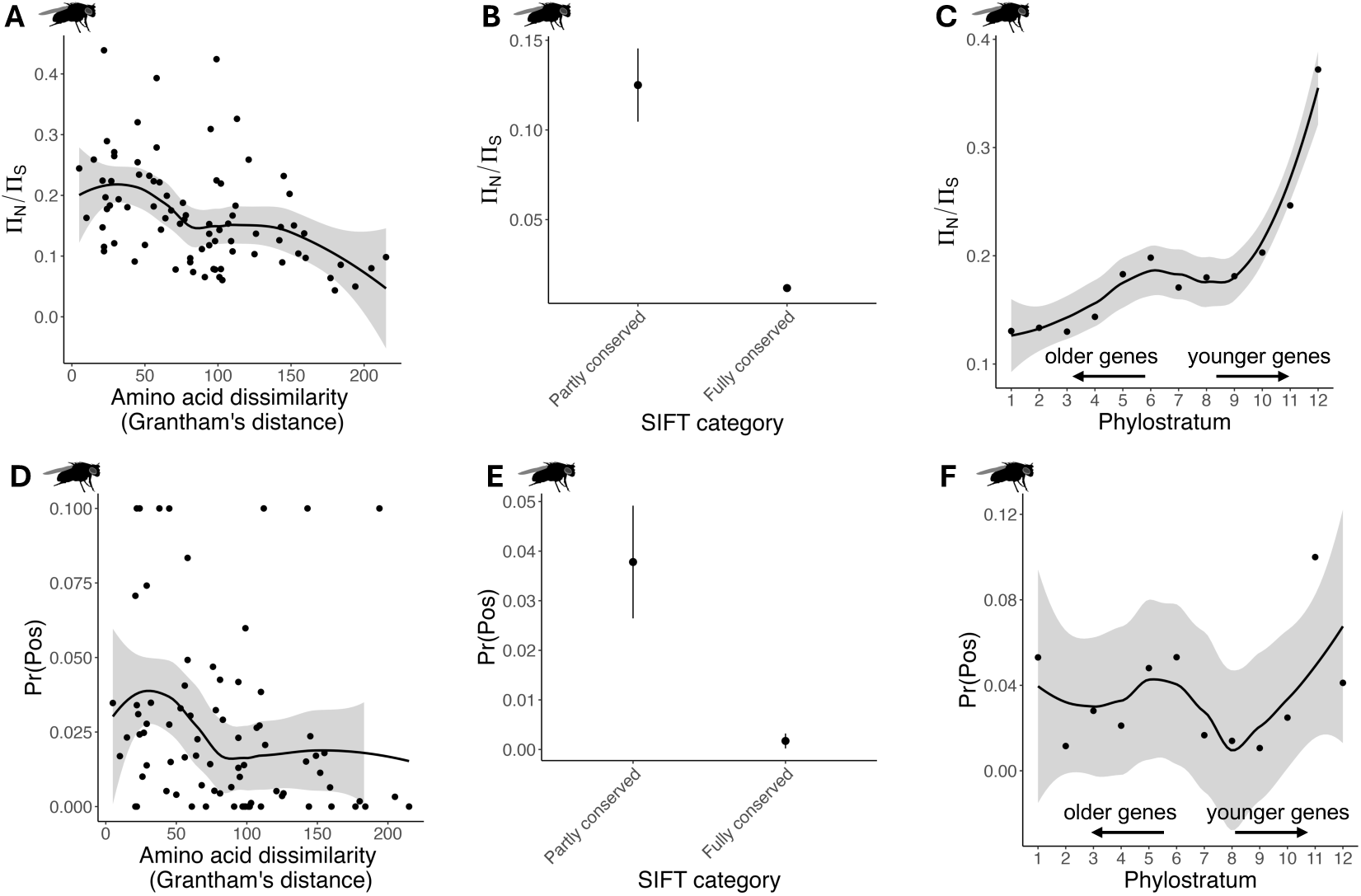
Proxies for effect size and ratios of autosomal nonsynonymous to synonymous nucleotide diversity in *D. melanogaster*. **A.** Point estimates of autosomal *π*_*N*_/*π*_*S*_, for polymorphisms affecting 81 pairs of amino acids, ranked by their Grantham’s distance, fitted with a loess regression curve. A significant negative rank correlation was observed (Spearman’s *ρ* = −0.461, *p* < 0.001). This was also found on the X (Spearman’s *ρ* = −0.446, *p* < 0.001; not shown). **B.** Same as A, but autosomal *π*_*N*_/*π*_*S*_was compared between “Partly conserved” and “Fully conserved” SIFT score categories (p<0.001, based on splitting the autosomal coding sequence into *n* bins with length equivalent to the X-linked coding sequence, and resampling autosomal estimates across these bins; points show the mean +/– one bootstrap standard error). **C.** Same as A, but autosomal *π*_*N*_/*π*_*S*_point estimates were compared between sites found in genes of varying phylostrata. Phylostratum 1 refers to “old” genes that are conserved across all cellular organisms, while phylostratum 12 represents “young” genes that are only found in Diptera. A significant positive correlation was observed between phylostratum and autosomal *π*_*N*_/*π*_*S*_ (Spearman’s *ρ* = 0.874, *p* < 0.001) corresponding to stronger purifying selection among older genes. This was also found for the X (Spearman’s *ρ* = 0.860, *p* < 0.001; not shown). **D.** Same as A, except shown for the proportion of sites under positive selection (“Pr(Pos)”) on autosomes as a function of Grantham’s distance (Spearman’s *ρ* = −0.359, *p* = 0.001). This correlation was also found on the X (Spearman’s *ρ* = −0.468, *p* < 0.001; not shown). **E.** Same as B, except shown for Pr(Pos) as a function of SIFT score (p<0.001). **F.** Same as C, except shown for Pr(Pos) as a function of gene age on autosomes (Spearman’s *ρ* = 0.056, *p* = 0.869) and the X (Spearman’s *ρ* = 0.504, *p* = 0.214; not shown).

Our second approach to validate the proxies was to examine the proportion of new mutations that are under positive selection (as estimated by fitting a DFE to unfolded site frequency spectra; see Materials and Methods) for mutations of different effect sizes. In Fisher’s geometric model, mutations with large scaled phenotypic effects are predicted to be less often under positive selection (Fisher 1930; Orr 1998)—and indeed, we found that large-effect mutations were less likely to be under positive selection for each of the three proxies (Fig. 3D-F), although the relationships are somewhat noisier than for *π*_*N*_/*π*_*S*_. We refrain from quantitatively estimating scaled phenotypic effects based on the proportion of mutations under positive selection (an assumption-laden exercise; see Materials and Methods), but this is in principle possible, as we show in the Supplementary Materials (Supplementary Fig. 16). In any case, the key point is that the *relative* values of our proxies tell us something about *relative* values of scaled phenotypic effects.

Having confirmed that our proxies for mutational effect size behave as expected, we correlated X-linked and autosomal rates of adaptation with each proxy (see Materials and Methods for details). We replicated the negative correlation between *ω*_*a*_ and effect size found previously in *D. melanogaster* (Fig. 4A-C) (Bergman and Eyre-Walker 2019; Moutinho et al. 2022), along with the strong signal of faster-X adaptation (Ávila et al. 2014; Campos et al. 2014; Charlesworth et al. 2018). However, we found no significant correlation between effect size and X-to-autosome ratios of *ω*_*a*_, whether using Grantham’s distances (Fig. 4D), SIFT scores (Fig. 4E) or gene age (Fig. 4F) as the metric of effect size. We repeated the analyses on 0- and 4-fold degenerate sites only (Supplementary Fig. 1), using folded site frequency spectra rather than unfolded site frequency spectra when estimating the DFE (Supplementary Fig. 2), using a more distant outgroup (*D. yakuba*, rather than *D. simulans*) (Supplementary Figs. 3-4), and splitting our data by sex-biased gene expression (Supplementary Figs. 5-7). We found no evidence of a relationship between X-to-autosome ratios of *ω*_*a*_and proxies for effect size, regardless of analysis strategy.

**Figure 4.**
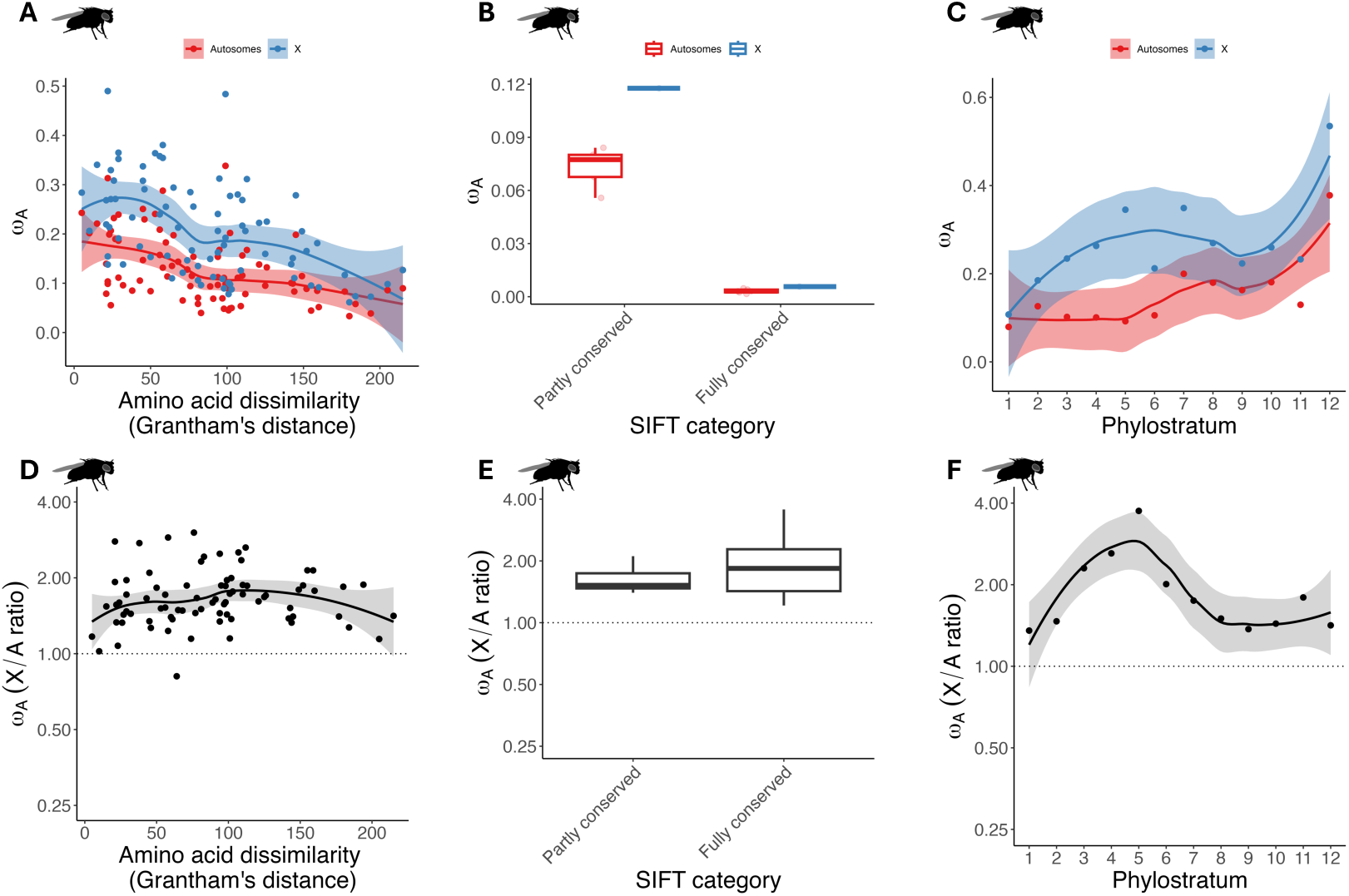
Proxies for effect size and relative rates of X-autosome adaptation in *D. melanogaster*. **A.** Point estimates of rates of adaptation, *ω*_*a*_, for nonsynonymous mutations affecting 81 pairs of amino acids, plotted against the Grantham’s distance between each amino-acid pair. **B.** Same as A, but for nonsynonymous mutations categorised as having “Partly conserved” or “Fully conserved” SIFT scores (boxplots were obtained by splitting the autosomal coding sequence into *n* bins with length equivalent to the X-linked coding sequence, and resampling estimates of *ω*_*a*_across these bins). **C.** Same as A, but for nonsynonymous mutations that fall in genes with different Phylostrata (smaller values indicate that genes are older). **D.** Correlation between X/A ratios of *ω*_*a*_ and Grantham’s distance (Spearman’s *ρ* = 0.129, *p* = 0.250), with X/A ratios plotted on a log2 scale. Negative values of *ω*_*a*_ were treated as zeroes and Infinite values were treated as 10^3^. **E.** X/A ratios of *ω*_*a*_ vs. SIFT score category (p=0.436, obtained by splitting the autosomal coding sequence into *n* bins with length equivalent to the X-linked coding sequence, resampling estimates of the X/A ratio of *ω*_*a*_ across these bins, and then calculating how many resampled estimates were larger in the “Fully conserved” than the “Partly conserved” SIFT category). **F.** Correlation between X/A ratios of *ω*_*a*_ and Phylostratum (Spearman’s *ρ* = −0.203, *p* = 0.528).

*D. melanogaster* is perhaps the ideal organism to test our hypothesis, due to its highly polymorphic coding sequence and its relatively large X chromosome. Still, we were curious to see if our results generalise across species. We therefore repeated our analyses (using Grantham’s distances and SIFT scores) in two species where faster-X adaptation has previously been reported (Kousathanas et al. 2014; Veeramah et al. 2014): the house mouse (*Mus musculus castaneus*), and humans. Although noisier, data from both species mirror those in *D. melanogaster* (Fig. 5A-D). We found a negative correlation between effect size and *π*_*N*_/*π*_*S*_ and *ω*_*a*_ (Supplementary Fig. 8), both on the X and autosomes, and we recovered the strong faster-X signals reported previously. However, we found little in the way of significant correlations between effect size and X/A ratios of *ω*_*a*_(Fig. 5A-D). One exception was a marginally significant (p=0.032) *negative* correlation between Grantham’s distance and X/A ratios of *ω*_*a*_ in *M. musculus*, but this was non-significant when using folded site frequency spectra to estimate the DFE (Supplementary Fig. 9). In general, we observed no significant relationships between effect size and X/A ratios of *ω*_*a*_, using different analysis strategies (Supplementary Figs. 9-15).

**Figure 5.**
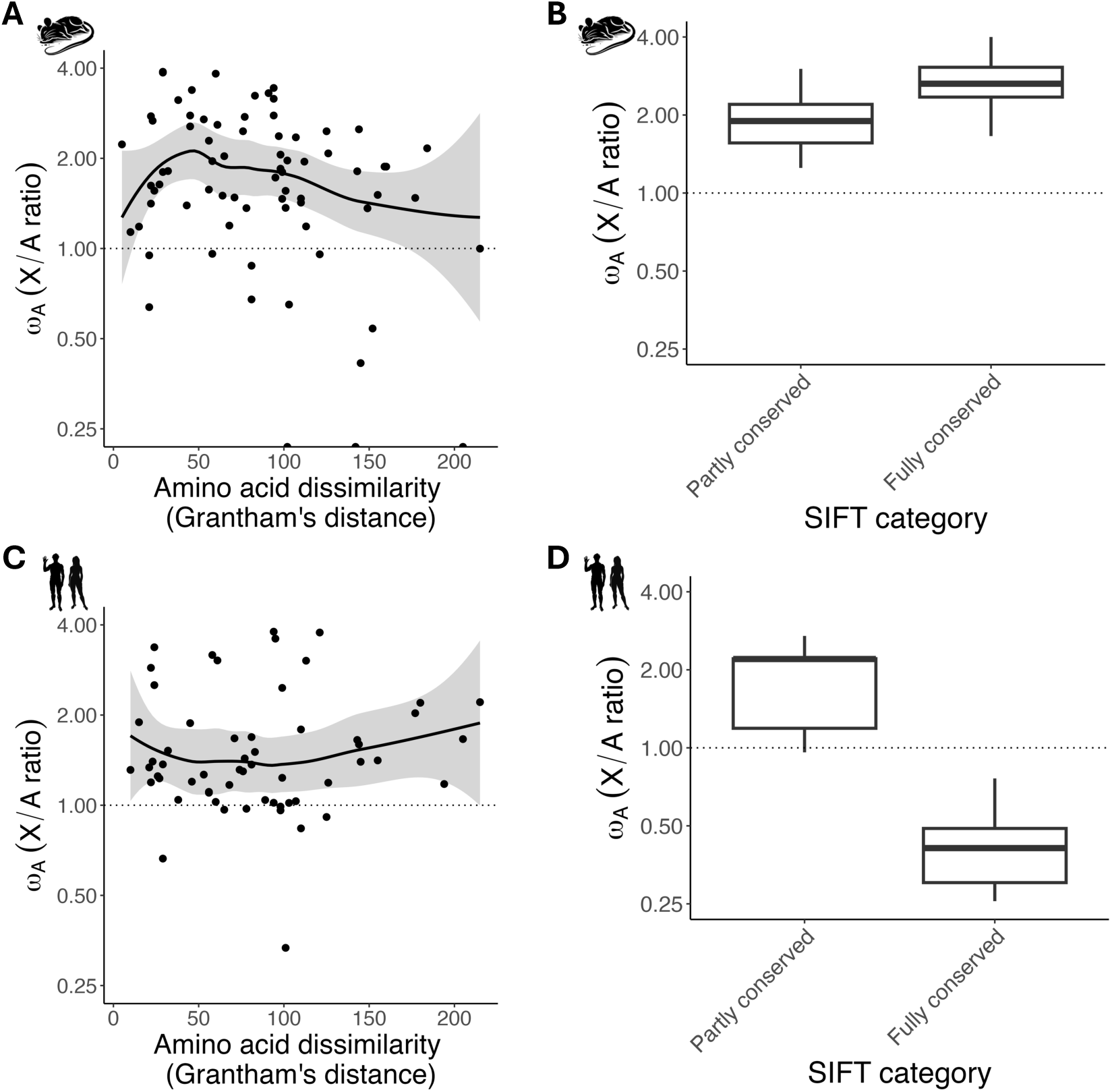
Proxies for effect size and relative rates of X-autosome adaptation in *M. musculus* and *H. sapiens*. **A.** Point estimates of X/A ratios of rates of adaptation, *ω*_*a*_, for nonsynonymous mutations affecting 81 pairs of amino acids, plotted against the Grantham’s distance between each amino-acid pair, in *M. musculus*. X/A ratios are plotted on a log2 scale. Negative values of *ω*_*a*_ were treated as zeroes and Infinite values were treated as 10^3^ (Spearman’s *ρ* = −0.239, *p* = 0.032). **B.** X/A ratios of *ω*_*a*_ vs. SIFT score category (p=0.136, obtained by splitting the autosomal coding sequence into *n* bins with length equivalent to the X-linked coding sequence, resampling estimates of the X/A ratio of *ω*_*a*_ across these bins, as shown by the boxplots, and calculating how many resampled estimates were larger in the “Fully conserved” than the “Partly conserved” SIFT category). **C.** Same as A., but for *H. sapiens* (Spearman’s *ρ* = 0.126, *p* = 0.260) **D.** Same as B., but for *H. sapiens* (p=0.879, using the same statistical approach as in B.).

## Discussion

Faster-X adaptation is a pervasive pattern in genome evolution (Meisel and Connallon 2013; McDonough et al. 2024). The most commonly invoked explanation—that new beneficial mutations are, on average, partially recessive (Charlesworth et al. 1987; Charlesworth et al. 2018)—is at odds with theories of dominance, which predict that new beneficial mutations should usually be partially dominant and/or overdominant for fitness (Kacser and Burns 1981; Manna et al. 2011). Sex differences in recombination, or sex differences in variance in reproductive success (Vicoso and Charlesworth 2009) may contribute to the effect, but they are at best partial explanations, especially in species where *N_eX_*/*N_eA_* ratios are equal to or smaller than ¾ (Carneiro et al. 2012; Hvilsom et al. 2012; Garrigan et al. 2014; Kousathanas et al. 2014; Veeramah et al. 2014; Pinharanda et al. 2019). Models of adaptation from standing genetic variation are even less promising, since they predict faster-autosome adaptation even when new beneficial mutations are partially recessive (Orr and Betancourt 2001). Here, we tested a new hypothesis: that faster-X adaptation arises due to large-effect mutations (McDonough et al. 2024). In McDonough et al. (2024)’s model, mutations with large scaled phenotypic effects can experience positive selection on the X even though equivalent mutations on autosomes experience balancing selection (and thus fail to fix) (Pamilo 1979). We tested the theory in three species (*D. melanogaster*, *M. musculus castaneus* and *H. sapiens*), using various proxies for mutational effect size (i.e., physico-chemical distances between amino-acid changing mutations, degrees of sequence conservation at nonsynonymous sites, and gene ages). Although we were able to replicate the faster-X effect in each species (Ávila et al. 2014; Campos et al. 2014; Kousathanas et al. 2014; Veeramah et al. 2014), we found no significant correlations between our proxies for scaled phenotypic effects and the extent of faster-X adaptation.

Why did we fail to detect positive correlations between our effect size proxies and the extent of faster-X adaptation? One possible explanation is that beneficial mutations have small “scaled phenotypic effects” (see Fig. 1B) and are typically recessive (we return to the issue of recessivity below). It is difficult to assess whether scaled phenotypic effects are likely to be large or small in general, as this depends on assumptions about the distribution of absolute phenotypic effects of mutations, mutational correlations among them, the number of independent traits, and the distance to the phenotypic optimum (Orr 1998). One argument against the idea that they are small is that most new mutations are deleterious. For example, mutation accumulation studies find that ∼90% of new mutations are deleterious (Bao et al. 2022), and fitting models of the DFE to nonsynonymous allele frequency spectra reveals a similarly high fraction of deleterious mutations (see Fig. 3D-F). This observation conflicts with infinitesimally-small scaled phenotypic effects, because half of new mutations should be deleterious and half should be beneficial in Fisher’s geometric model when this is the case. The low proportion of beneficial mutations instead suggests that scaled phenotypic effects will often be large (see Supplementary Fig. 16).

Another possible contributor to our negative results is the difficulty of accurately estimating scaled phenotypic effects. We have relied on proxies, which are intended to capture variation in absolute phenotypic effects (i.e., Grantham’s distances) or in the distance to the optimum (i.e., SIFT scores, gene age). These proxies are imperfect. For example, different components of scaled phenotypic effects might trade-off with one another (e.g., mutations in older genes of *D. melanogaster* have smaller Grantham’s distances and are more pleiotropic than younger genes (Moutinho et al. 2022; Martin and Tate 2025)), which might dampen any genuine signal. One possible solution could be to develop a composite metric that incorporates absolute phenotypic effects (e.g., via Grantham’s distances), distance to the optimum (e.g., via gene age) and the number of independent traits (e.g., some measure of gene pleiotropy). Such a metric might then better capture scaled phenotypic effects and improve the power of the test. A related problem is that our proxies do not necessarily capture the full range of scaled phenotypic effects. For example, mutations with the smallest Grantham’s distances might still have large scaled phenotypic effects (see Supplementary Fig. 16), which would then (under McDonough et al.’s theory) lead to faster-X adaptation across the range of Grantham’s distances. But even if this is true, an observation that requires explanation is the absence of a correlation between effect size and faster-X, which fits poorly with McDonough et al.’s theory, assuming semi-dominant phenotypic effects (dark green line in Fig. 1B). Might some other distribution of phenotypic dominance generate a relatively constant faster-X across effect size classes (e.g., light green line for *v* = 0.25 in Fig. 1B)?

With regards to data on dominance, there is a well-established observation that new deleterious mutations are partially recessive (Simmons and Crow 1977; Charlesworth 1979; Orr 1991; Agrawal and Whitlock 2011; Manna et al. 2011), but data on phenotypic dominance, or on the dominance of beneficial mutations, is much sparser (Orr 2010; Billiard et al. 2021; Di and Lohmueller 2024). In the case of mutations conferring pesticide resistance in *Drosophila*, Bourguet & Raymond (1998) found that most mutations generate enough enzyme activity to be dominant for survival, while Ronfort & Glémin (2013) found a similar pattern when looking at quantitative trait loci affecting domestication in outcrossing plants. However, both estimates suffer from the well-known bias towards the fixation of dominant mutations in outcrossing diploids (i.e., Haldane’s sieve (Haldane 1927; Orr and Betancourt 2001)). In the same study by Ronfort & Glémin (2013), phenotypic dominance was also estimated among self-fertilising plants, where Haldane’s sieve should be minimal (Charlesworth 1992). There, they found a distribution shifted towards partially recessive effects. As these authors point out, however, it is not clear how domestication is related to total fitness, so to extrapolate from their estimates to dominance for fitness requires some caution.

Further direct estimates of the dominance of beneficial mutations come from evolution experiments in *S. cerevisiae*. In the case of haploid yeast lineages adapting to the antifungal fluconazole, roughly half of the ∼30 adaptive mutations identified were recessive when they were expressed in diploids, while the other half were dominant (Anderson et al. 2004). For a dozen mutations conferring adaptation to nystatin, all were recessive (Gerstein et al. 2014). However, the candidate mutations identified in these experiments were often loss-of-function mutations affecting a single genetic pathway. Such loss-of-function mutations are unlikely to represent the genetic basis of adaptation across a broad set of environments, and such mutations should contribute minimally to signals of molecular adaptation based on MK tests. Accordingly, further experiments conducted in different environments have found that candidate adaptive mutations often exhibit dominance or overdominance for fitness (Sellis et al. 2016; Marad et al. 2018; Aggeli et al. 2022). Overall, direct data on the dominance of new beneficial mutations remains exceptionally patchy, and there is a pressing need for larger-scale data in a range of contexts of selection to help parameterise models of adaptation (Orr 2010). “Deep mutational scanning” approaches (Padhy et al. 2023; Allen et al. 2026), in which a large number of mutations are generated (in haploid or diploid state) and their fitness effects are inferred from allele frequency trajectories among competing clones offer promise in this regard.

Indirect inferences into the dominance of beneficial mutations have also been made by contrasting X and autosome adaptation among genes with male-biased vs. female-biased fitness effects. In Charlesworth et al.’s classical model of faster-X adaptation (Charlesworth et al. 2018), mutations that are only expressed in females are expected to evolve equally fast on autosomes and the X, while mutations that are only expressed in males are expected to show a strong faster-X effect. Evidence that genes with male-biased expression experience stronger faster-X adaptation is mixed: it is observed in *D. melanogaster* (Campos et al. 2018) and *M. musculus* (Kousathanas et al. 2014), but not in *H. sapiens* (Veeramah et al. 2014) or *H. melpomene* (Pinharanda et al. 2019) (a ZW system, in which the prediction applies to female-biased genes). The weak relationship between faster-X adaptation and sex-biased gene expression is unsurprising, given that sex-biased expression is (at best) an imperfect proxy for sex-specific fitness effects. For example, sex-biased gene expression is positively related to metrics of sex-differential selection in some datasets (Connallon and Clark 2011), while other datasets find no (Ming et al. 2025), mixed (Grieshop et al. 2025) or negative (Ruzicka et al. 2019) associations. Still, establishing whether faster-X adaptation typically occurs for mutations with male-limited effects will be a valuable avenue for further work, given that McDonough et al. (2024)’s model only predicts faster-X under male-limited expression if new mutations show recessivity at the trait level.

Faster-X models typically consider scenarios where adaptation is limited by the supply of new mutations, but it is plausible that adaptation uses pre-existing genetic variation (Barrett and Schluter 2008). But from the perspective of faster-X, adaptation from standing genetic variation is even less conducive to faster-X than mutation-limited models (Orr and Betancourt 2001; Connallon et al. 2012). For example, Orr & Betancourt (2001) considered a population with standing genetic variation maintained at a balance between recurrent mutation and purifying selection. The population then experiences a sudden shift in environmental conditions, such that deleterious alleles become beneficial while retaining the same dominance coefficient in the old and new environment. In this model, X-linked substitution rates are invariably lower than autosomal substitution rates. Not only do X-linked mutations suffer from reduced mutational input (as they do in mutation-limited models), but they are further hampered by their lower initial frequencies at the time of the environmental shift. Still, there is currently no thorough theoretical exploration of X and autosomal substitution rates in scenarios in which adaptation involves the simultaneous segregation of many mutations (e.g., polygenic models of stabilising selection at a fixed optimum (Barton and Turelli 1989), stabilising selection of a randomly fluctuating optimum (Bertram and Shafiei 2025), adaptation to a sudden shift in optimum (Hayward and Sella 2022)). Such theory would be extremely valuable to identify conditions, if any, that facilitate faster-X adaptation when adaptation is not mutation-limited (see a recent simulation study (Muralidhar and Coop 2023) for a step in this direction). Such theory might also help us to interpret the apparent enrichment of hard selective sweeps on the X chromosome and soft selective sweeps on autosomes (Harris and Garud 2023).

Another important theoretical assumption in models of faster-X is that the fitness effects of beneficial mutations in hemizygous males are equivalent to those in homozygous females. This assumption is well supported by data from the UK Biobank in humans (Sidorenko et al. 2019), in which the phenotypic effects of X-linked hemizygous mutations in males are typically equivalent to X-linked homozygous effects in females. However, there is wide variation in the mechanisms and the extent of dosage compensation across species (Cecalev et al. 2024), which may influence the relative fitness effects of mutations between females and males on the X chromosome. Empirical work investigating how variation in dosage compensation is related to the phenotypic and fitness effects of mutations (and thus, presumably, to patterns of X-linked variation and selection) is therefore needed to solidify this assumption. Nonetheless, the fact that faster-X is observed in a variety of dosage compensation systems—from global upregulation of X-linked gene expression in male *D. melanogaster*, to random inactivation of most X-linked genes in female placental mammals—suggests that the explanation for faster-X is robust to variation in dosage compensation mechanisms.

Finally, our inferences about rates of adaptation rely on the MK test, which is a robust (Booker 2020) but not infallible tool. On the one hand, some potential biases of the MK test (e.g., the assumption that the DFE is stable on the timescale of divergence) are alleviated by X-autosome contrasts, since such biases should apply equally to both chromosome types. On the other hand, X-autosome contrasts introduce new complications. For example, a standard assumption in current implementations of the MK test is that segregating functional polymorphisms are either neutral, unconditionally deleterious or unconditionally beneficial (Eyre-Walker and Keightley 2009; Galtier 2016). Yet, segregating functional polymorphisms may also be overdominant (Sellis et al. 2011; McDonough and Connallon 2023; McDonough et al. 2024; Ruzicka et al. 2025). If overdominant polymorphisms are enriched on autosomes, as predicted by theory (Pamilo 1979), this “hidden” overdominance could exacerbate signals of faster-X adaptation. Whether overdominant polymorphisms are in fact enriched on autosomes remains to be verified: to our knowledge, X-autosome contrasts have only been performed among candidates for long-term balancing in humans, and such candidates loci are equally abundant on both chromosome types (see Supplementary Materials in (Leffler et al. 2013)). Whether the lack of enrichment applies to targets of short-term balancing selection (Soni et al. 2022), and whether this appreciably distorts MK tests, remains to be investigated.

## Materials and Methods

### Divergence data

Our three focal species were *D. melanogaster*, *M. musculus*, and *H. sapiens*. The relevant reference assemblies were number 5 for *D. melanogaster* (GCF_000001215.4), GRCm38 for *M. musculus* (GCF_000001635.20), and GRCh38 for *H. sapiens* (GRCh38.p14). For each focal species, we obtained DNA and protein sequences for all transcripts using gffread (Pertea and Pertea 2020), a tool that takes the gff and fasta files for each species’ reference as input and considers the strand and reading frame of each transcript. For each gene, we removed all but the longest transcript, using custom bash scripts. To identify substitutions, we used two outgroups for each lineage: *D. simulans* and *D. yakuba*, which have diverged from *D. melanogaster* 1–4 million years ago (MYA), and 2–9 MYA, respectively (Obbard et al. 2012; Suvorov et al. 2022); *M. spretus* and *M. pahari*, which have diverged from *M. musculus* 1–2 MYA and 5–7 MYA, respectively (Suzuki et al. 2004; Thybert et al. 2018); and *Pan troglodytes* and *Gorilla gorilla*, which have diverged from humans 7–8 MYA, and 11–17 MYA, respectively (Langergraber et al. 2012). In each lineage, we identified 1:1:1 orthologs using orthofinder (Emms and Kelly 2019), resulting in 11,716 orthogroups for the *Drosophila* lineage, 14,862 for the mouse lineage and 14,949 for the primate lineage. We then aligned each orthogroup in turn, using the codon-aware multiple sequence aligner MACSE (Ranwez et al. 2011). We recovered the position of every site in terms of the coordinates of the focal species’ reference genome, using custom bash scripts. We only considered substitutions (e.g., between *D. melanogaster* and *D. simulans*) that were non-polymorphic in the focal species (e.g., in *D. melanogaster*).

### Polymorphism data

For *D. melanogaster*, we used single nucleotide polymorphism (SNP) data from a Zambian population sample (DPGP3), downloaded from www.johnpool.net/genomes.html. The sample consists of 197 haploid genome sequences from *D. melanogaster’*s ancestral distribution range, and is minimally affected by the bottlenecks accompanying *D. melanogaster*’s expansion out of Africa (Arguello et al. 2019). Details of the SNP calling pipeline can be found in (Lack et al. 2016). For each major chromosome arm, the multiple-alignment fasta files were converted to vcf using snp-sites (Page et al. 2016), and coding (CDS) regions were extracted using bedtools (Quinlan and Hall 2010). We obtained allele counts for polymorphic sites using vcftools (Danecek et al. 2011), retaining only biallelic sites, sites with call rate >80% across individuals, and sites where at least one of the polymorphic alleles matches the transcribed allele in the reference assembly, using custom R scripts (RStudio Team 2020). Because alleles in the polymorphism dataset are reported with respect to the forward strand (even if transcription takes place on the reverse strand), we converted alleles to their complement whenever the site was transcribed from the reverse strand (e.g., if a C/T polymorphism was transcribed from the reverse strand, we converted it to a G/A polymorphism).

For *M. musculus*, we used polymorphism data from a sample of 10 diploid individuals of the sub-species *M. musculus castaneus*, whose ancestral distribution range is in India and whose *N_e_* is inferred to be larger than *M. musculus domesticus* (Halligan et al. 2010). Reads had previously been mapped to the GRCm38 reference assembly (see (Harr et al. 2016) for details) and were downloaded from wwwuser.gwdguser.de/~evolbio/evolgen/wildmouse/vcf/. We applied the same filters as in *D. melanogaster*, and we further removed pseudo-autosomal regions (positions 90,745,845–91,644,698 and 169,969,759–170,931,299; per <u>/GCF_000001635.20_GRCm38_assembly_structure/genomic_regions_definitions.txt</u>). For *H. sapiens*, we used polymorphisms from the 1000 genomes project (The 1000 Genomes Project Consortium 2015), aligned to the GRCh38 assembly, and downloaded from www.cog-genomics.org/plink/2.0/resources. We initially converted pgen files to vcf files using plink2 (Chang et al. 2015), and then proceeded as elsewhere, removing pseudo-autosomal regions as well (positions 10,001–2,781,479 and 155,701,383–156,030,895; per GCA_000001405.29_GRCh38.p14_assembly_structure/genomic_regions_definitions.txt).

We used parsimony to infer ancestral allele states at the node of the tree leading to each focal species. Specifically, we inferred an allele to be ancestral if both outgroup species (e.g., *D. simulans* and *D. yakuba*) carried the allele; the alternative allele was then assigned as the derived allele. In cases where the focal species carried an allele that was not present in the two outgroup species, we removed the polymorphism from further analysis. In cases where the more distant outgroup (e.g., *D. yakuba*) was used to estimate divergence, we inferred ancestral allele state based on this outgroup only.

### Scaled phenotypic effects and their proxies

Before describing how we estimated the effect size of mutations, it is worth clarifying what kind of effect size we are aiming to measure, since there are many possible kinds in population genetics. For our purposes, the relevant effect size is a mutation’s “scaled phenotypic effect” (*x*), which is formally defined as 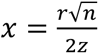, where *r* is an absolute phenotypic effect (summed across a set of independent traits), *n* is the number of independent traits, and *z* is the distance from the phenotypic optimum (Orr 1998). Thus, mutations can have large “*x*” if they have large absolute phenotypic effects (i.e., large values of *r*), or if they are close to the optimum (i.e., small values of *z*), as illustrated in the left vs. right panels of Fig. 1A. Put simply, our first metric of effect size (Grantham’s distances; see below) is intended to capture variation in absolute effect size (*r*), while our two other metrics (SIFT score and gene age; see below) are intended to capture variation in distance to the phenotypic optimum (*z*).

Values of *x* are very difficult to estimate empirically. Estimating *x* requires making assumptions about the distribution of absolute phenotypic effects, the number of independent traits, and mutational correlations among them. Moreover, if we model adaptation as an “adaptive walk”, populations closer to the optimum should tend to fix mutations with smaller *x* than populations far from the optimum (Orr 1998; Moutinho et al. 2022), so that the value of *x* will also depend on assumptions about which step in the adaptive walk the population is in. Despite these limitations, proxies can still provide information about the *relative* values of *x* among different classes of mutation. Moreover, we can validate the proxies by asking whether large-effect mutations, thus inferred, show properties that are consistent with those predicted for large-*x* mutations in Fisher’s geometric model. For example, a long-standing result (Fisher 1930) is that the proportion of mutations under purifying selection should increase as *x* increases. We tested this by asking whether large-effect mutations, inferred from our proxies, were more likely to be under purifying selection than small-effect mutations, using *π*_*N*_/*π*_*S*_, a common measure of the efficiency of purifying selection (Castellano et al. 2018). Another prediction is that the proportion of mutations under positive selection should decrease as *x* increases (Fisher 1930). The proportion of mutations under positive selection was estimated from fitting models of the distributions of fitness effects (DFE) to nonsynonymous and synonymous polymorphism data, using the software grapes (Galtier 2016) (see details below). Finally, by way of illustration, we present analytical predictions that relate the average value of *x* to the estimated proportion of mutations under positive selection (see Supplementary Fig. 16).

### Proxies for mutational effect size

We used three metrics as proxies for scaled phenotypic effects. First, we partitioned nonsynonymous mutations by dissimilarity using Grantham’s distances, downloaded from gist.github.com/danielecook/501f03650bca6a3db31ff3af2d413d2a#file-grantham-tsv. The Grantham’s distance between a pair of amino acids is a composite measure of the dissimilarity in polarity, volume, and composition between them (Grantham 1974). In other words, it quantifies whether a mutation converts one amino acid (e.g., Isoleucine) to a very similar amino-acid (e.g., Leucine) or to a very dissimilar amino acid (e.g., Serine). Thus, it should capture variation in the absolute phenotypic effects of mutations. In practice, we predicted the effects of all possible coding mutations at every site in each focal species using degenotate (Mirchandani et al. 2024). Mutations (polymorphisms and substitutions) were then assigned a Grantham’s distance using custom R scripts.

Second, we partitioned nonsynonymous mutations by their “Sorting Intolerant from Tolerant for Genomes” (SIFT) score (Ng and Henikoff 2001). SIFT is an algorithm to predict the functional consequences of mutations based on a multiple sequence alignment with related species. Our rationale is that a highly conserved site is likely to be close to its optimum, such that a mutation affecting a highly conserved site should have a larger scaled phenotypic effect than a weakly conserved site (given that scaled phenotypic effects are inversely proportional to the distance to the optimum in Fisher’s geometric model). SIFT scores have previously been used to partition putatively beneficial mutations into those of small and large effect (Chen et al. 2022). In practice, we used SIFT4G (Vaser et al. 2016) to calculate SIFT scores for each possible mutation at every coding site, using publicly available SIFT files (BDGP5.74, GRCm38.74, GRCh38.74, respectively, downloaded from https://sift.bii.a-star.edu.sg/sift4g/public/) and reference genome files (converted to vcf) as input. SIFT scores vary between 0 and 1 and are highly skewed towards 0. We assigned a SIFT score of 1 to reference alleles, and categorised large-effect (“partly conserved”) mutations as those where the difference in SIFT scores between two alleles was >0.9999, and small-effect mutations (“fully conserved”) as those where the difference in SIFT scores was 0.0499-0.9999.

Finally, we used a phylostratigraphic approach to classify nonsynonymous mutations. Mutations were binned by gene age, into “old” genes that share homologues in distantly related species and “young” genes that only share homologues in closely related species. The rationale is that older genes should be closer to their optimum, such that mutations in older genes should have larger scaled phenotypic effects. To classify genes by age in *D. melanogaster*, we used the 12 phylostrata defined by (Domazet-Lošo et al. 2017).

### Rates of adaptation

We used MK tests to estimate rates of adaptation (McDonald and Kreitman 1991). MK tests compare polymorphism (*P*_*s*_) and divergence (*D*_*s*_) at putatively neutral sites, and polymorphism (*P*_*N*_) and divergence (*D*_*N*_) at putatively functional sites. If all new mutations at putatively functional sites are either neutral or strongly deleterious, then 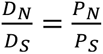. If some putatively functional mutations are adaptive, the number of excess substitutions due to positive selection can be estimated as 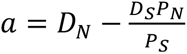 (Smith and Eyre-Walker 2002). In the absence of segregating non-neutral mutations, the proportion of putatively functional substitutions that are adaptive can then be estimated as 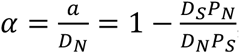, and the rate of adaptation (scaled by the rate of neutral substitution) is 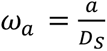, (Galtier 2016). We used *ω*_*a*_ as our focal metric of adaptation because it is a rate, rather than a proportion, and thus more directly relevant to the theory (similar results were found using *α* instead; not shown).

In practice, mildly deleterious mutations often segregate at putatively functional sites, which will downwardly bias estimates of *α* and *ω*_*a*_ (Keightley and Eyre-Walker 2007; Eyre-Walker and Keightley 2009). To correct for this, we fitted models of the DFE among putatively functional polymorphisms, using grapes (Galtier 2016). Specifically, we fitted the “GammaExpo” model, where segregating deleterious mutations are assumed to follow a gamma distribution (whose shape and scale parameters are estimated by grapes) and segregating beneficial mutations follow an exponential distribution. This model accurately estimates rates of adaptation (Booker 2020) and provides the best fit to unfolded allele frequency spectra in most species (Rousselle et al. 2020). To guard against mis-estimation of ancestral allele states, we repeated analyses on the folded allele frequency spectrum, fitting the “GammaZero” model, which considers that segregating beneficial mutations are absent and segregating deleterious mutations follow a gamma distribution. This model also provides an excellent fit in most species (Rousselle et al. 2020). Because DFE estimation in grapes does not handle large samples of sequences well, we down-sampled the allele frequency spectrum to a maximum of 20 sequences. We did this by splitting the full site frequency spectrum into 20 bins and multiplying counts in each bin by a factor 20/n, where n is the number of sequences in the sample (e.g., 20/197 for *D. melanogaster* polymorphism data). 20 sequences is sufficient for accurate inferences of the DFE and *ω*_*a*_(Charlesworth and Eyre-Walker 2008), provided that the number of segregating sites per unit of analysis is large (on the order of 10^3^-10^4^ SNPs, per (Andersson et al. 2023)).

Grapes requires a mutational target size as input. To estimate mutational target size, we considered the base composition of coding sequences and transition-transversion (ts:tv) bias. Specifically, given that each base occurs at a fraction *f* in a focal species’ coding sequence, we assumed the probability of a transversion to be 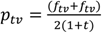 and the probability of a transition to be 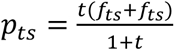, where *f*_*tv*_ and *f*_*ts*_ are pairs of bases undergoing transversions and transitions, respectively, and *t* is the ts:tv ratio of mutation *counts*. Based on mutation accumulation data, we used *t* = 1 for *D. melanogaster* (Keightley et al. 2009) and *t* = 2 for *Mus* and humans (Kong et al. 2012; Lynch et al. 2023), corresponding to 2-fold and 4-fold higher transition *rates*, respectively. We estimated mutational target size for a given site class by summing all possible mutations of that class across a species’ coding sequence, weighted by its mutation probability (given overall genomic base composition and relevant ts:tv ratios).

In many cases, mutations in a particular amino acid pair (e.g., Alanine to Valine) originate from a few (often just one) kind of mutation (e.g. C↔T), which can cause problems (when comparing across mutational classes) if there is codon usage bias (Singh et al. 2008; Vicoso et al. 2008) or GC-biased gene conversion (which typically favour G or C alleles (Jackson et al. 2017; Rousselle et al. 2019)). To account for this, we followed the approach of Bergman & Eyre-Walker (2019). Specifically, for a given amino acid pair, we used synonymous sites that are separated by the same mutation type as the neutral baseline, such that all relevant inputs for grapes consisted of that mutation type only (e.g., C↔T mutations only, for the Alanine to Valine amino acid pair). When an amino acid pair was separated by more than one mutation type (e.g., some mixture of C↔T and C↔G), we weighted synonymous sites by the relative frequency of each mutation type (e.g., if a given amino acid pair was predicted to originate via 50% C↔T and 50% C↔G mutations, the synonymous site frequency spectrum was weighted 50% to C↔T mutations and 50% to C↔G mutations).

### Recombination rates

In *D. melanogaster*, recombination rate estimates were obtained from Comeron et al. (2012), downloaded from https://comeron.lab.uiowa.edu/recombination-rates. The estimates are provided in 100Kb windows, which can result in noisy estimates of *ω*_*a*_if there are few polymorphisms in each window. To minimise this effect, we averaged recombination rates across 10 non-overlapping windows, and assigned each site to its relevant 1Mb window. Since recombination rates were estimated in females and there is no recombination in male *Drosophila*, we multiplied recombination rate estimates by 2/3 for the X and by ½ for autosomes. These corrections adjust for the fact that two-thirds of X-linked genes and one-half of autosomal genes are carried by females.

### Sex-biased gene expression

The sex-specific fitness effects of mutations should correlate with X/autosome ratios of adaptive substitution rates (Charlesworth et al. 2018), and sex-biased expression is often used as a proxy for the sex-specific fitness effects (whether this assumption is justified is something we return to in the Discussion). We therefore examined rates of adaptation among genes with different levels of sex-biased expression. We used expression data that has been pre-processed by Fraïsse et al. (2019). Briefly, the expression data consist of numbers of reads per kilobase per million reads (RPKM), obtained from Flybase (“gene_rpkm_report_fb_2017_04.tsv.gz”). These counts have been quantile-normalised between males and females, and between tissues for a given developmental stage. We then estimated the sex-bias ratio as Fraïsse et al. (2019) did, by estimating the mean of reproductive tissue samples in females (ovary samples among 4-day-old mated and virgin females) and males (accessory gland and testis samples among 4-day-old males), and taking the log2 ratio of the male-to-female values. Male-biased, female-biased and unbiased genes were classified as genes with a log2 ratio greater than 1, smaller than –1, and between –1 and 1, respectively.

## Data availability statement

All data and code necessary for confirming the conclusions of the article can be found at https://github.com/filipluca/FasterX_and_mutation_size

## Acknowledgments

We thank Tim Connallon for comments on a draft of this manuscript and for assistance with the theory presented in the Supplementary Materials.

