## Supplementary Materials for "Is faster-X adaptation due to large-effect mutations? An empirical test of a new theory"

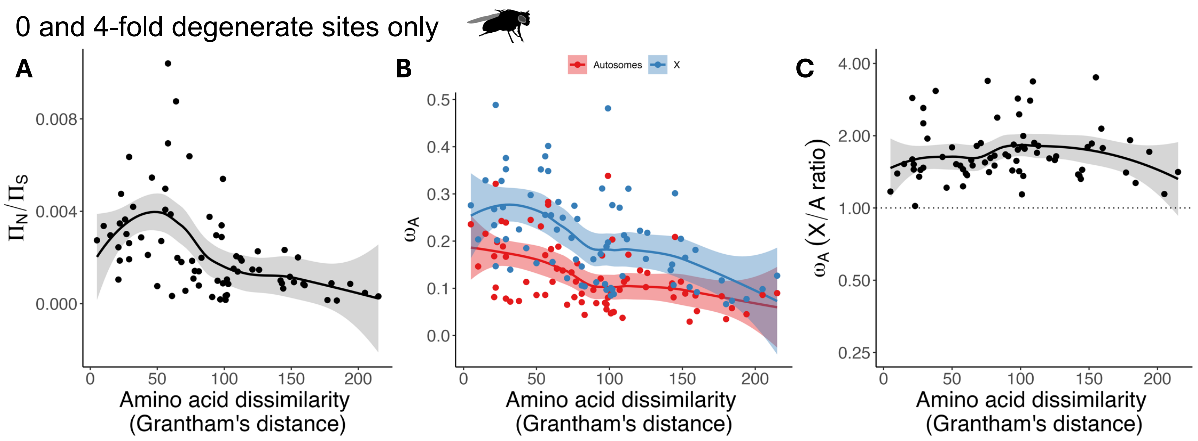


**Supplementary Figure 1. Grantham's distances, ratios of autosomal nonsynonymous to synonymous nucleotide diversity, and relative rates of X-autosome adaptation in *D. melanogaster*, for 0 and 4-fold degenerate sites. A.** Autosomal $\pi_0/\pi_4$ estimates, for polymorphisms affecting each pair of amino acid, ranked by Grantham's distance, fitted with a loess regression curve. A significant negative rank correlation was observed (Spearman's $\rho$=-0.577, $p<0.001$). **B.** Rates of adaptation, $\omega_a$, for nonsynonymous mutations affecting each pair of amino acids, plotted against the Grantham's distance between each amino-acid pair. **C.** Correlation between X/A ratios of $\omega_a$ and Grantham's distance (Spearman's $\rho = 0.087, p = 0.463$), with X/A ratios plotted on a log2 scale. Negative values of $\omega_a$ were treated as zeroes and Infinite values were treated as $10^3$.

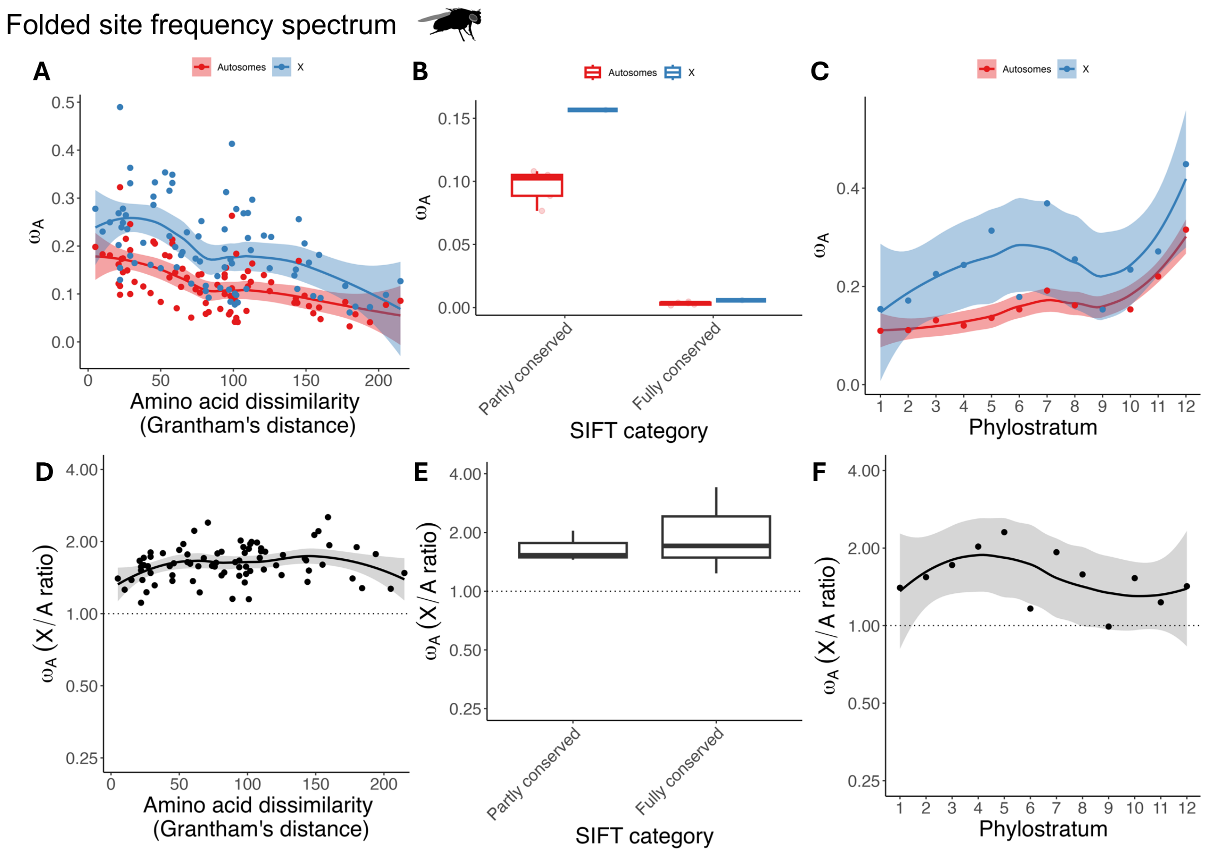


**Supplementary Figure 2. Proxies for effect size and relative rates of X-autosome adaptation in *D. melanogaster*, using the "GammaZero" model for fitting the DFE to the folded site frequency spectrum. A.** Rates of adaptation, $\omega_a$, for nonsynonymous mutations affecting 81 pairs of amino acids, plotted against the Grantham's distance between each amino-acid pair. **B.** Same as A, but for nonsynonymous mutations categorised as having "Partly conserved" or "Fully conserved" SIFT scores. **C.** Same as A, but nonsynonymous mutations that fall in genes with different Phylostrata (smaller values indicate that genes are older). **D.** Correlation between X/A ratios of $\omega_a$ and Grantham's distance (Spearman's $\rho$=0.227, p=0.041), with X/A ratios plotted on a log2 scale. Negative values of $\omega_a$ were treated as zeroes and Infinite values were treated as $10^3$. **E.** X/A ratios of $\omega_a$ vs. SIFT score category (p=0.466, obtained by splitting the autosomal coding sequence into *n* bins with length equivalent to the X-linked coding sequence, resampling estimates of the X/A ratio of $\omega_a$ across these bins, and then calculating how many resampled estimates were larger in the "Fully conserved" than the "Partly conserved" SIFT category). **F.** Correlation between X/A ratios of $\omega_a$ and Phylostratum (Spearman's $\rho$=-0.350, p=0.266).

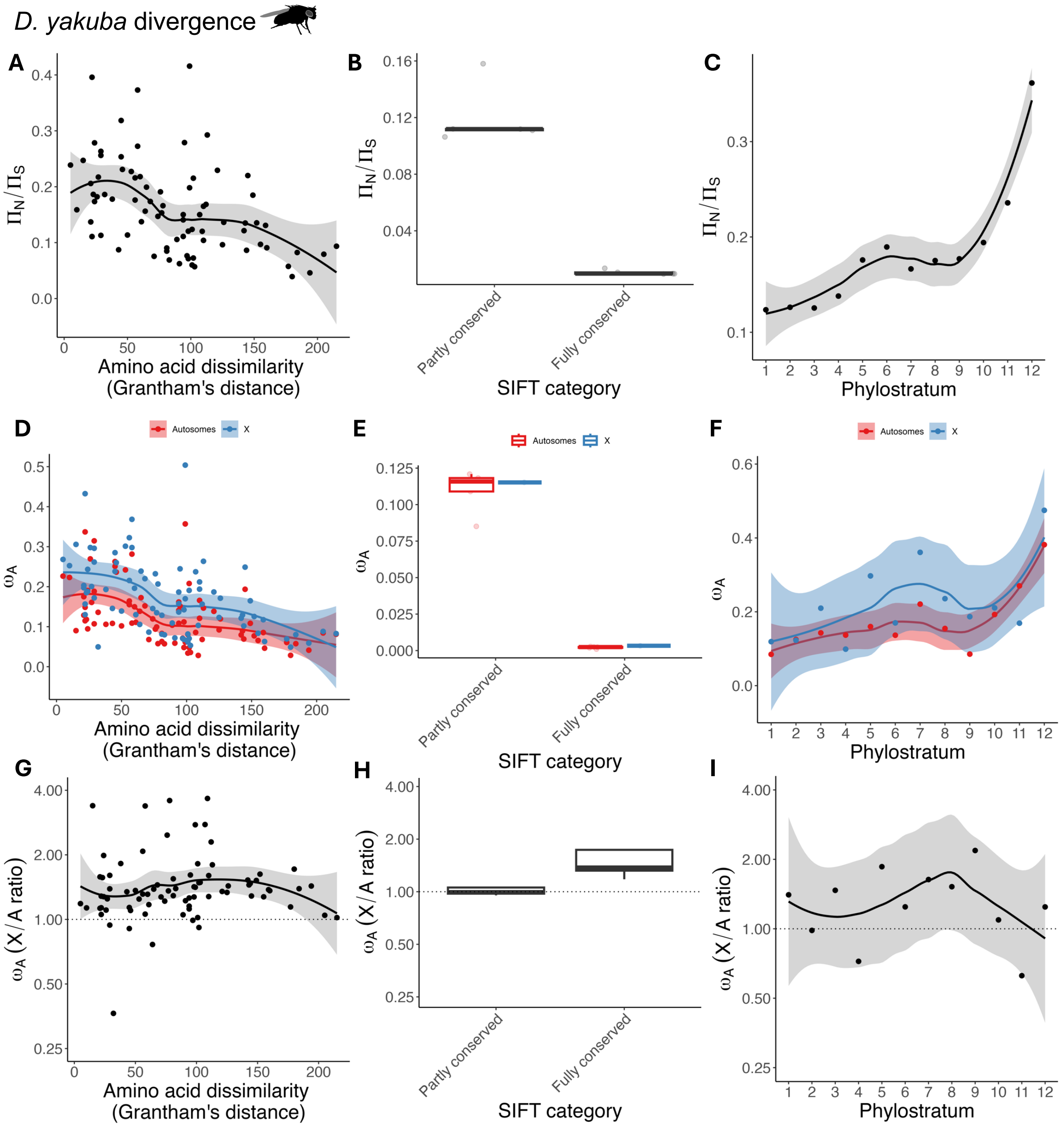


**Supplementary Figure 3. Proxies for effect size, ratios of autosomal nonsynonymous to synonymous nucleotide diversity, and relative rates of X-autosome adaptation in *D. melanogaster*, calculated from divergence to *D. yakuba*. A.** Autosomal $\pi_N/\pi_S$ estimates, for polymorphisms affecting 81 pairs of amino acids, ranked by their Grantham's distance, fitted with a loess regression curve. A significant negative rank correlation was observed (Spearman's $\rho$=-0.476, $p<0.001$). **B.** Same as A, but autosomal $\pi_N/\pi_S$ is compared between "Partly conserved" and "Fully conserved" SIFT score categories ($p<0.001$, based on splitting the autosomal coding sequence into n bins with length equivalent to the X-linked coding sequence, and resampling autosomal estimates across these bins). **C.** Same as A., but autosomal $\pi_N/\pi_S$ is compared between sites found in genes of varying phylostrata. Here, phylostratum 1 refers to "old" genes that are conserved across all cellular organisms, while phylostratum 12 represents "young" genes that are only found in Diptera. A significant positive correlation was observed between phylostratum and autosomal $\pi_N/\pi_S$ (Spearman's $\rho$=0.916, $p<0.001$), corresponding to stronger purifying selection among older genes. **D.** Rates of adaptation, $\omega_a$, for nonsynonymous mutations affecting 81 pairs of amino acids, plotted against the Grantham's distance between each amino-acid pair. **E.** Same as D, but for nonsynonymous mutations categorised as having "Partly conserved" or "Fully conserved" SIFT scores. **F.** Same as D, but nonsynonymous mutations that fall in genes with different Phylostrata (smaller values indicate that genes are older). **G.** Correlation between X/A ratios of $\omega_a$ and Grantham's distance (Spearman's $\rho$=0.185, p=0.133), with X/A ratios plotted on a log2 scale. Negative values of $\omega_a$ were treated as zeroes and Infinite values were treated as $10^3$. **H.** X/A ratios of $\omega_a$ vs. SIFT score category (p=0.065, obtained by splitting the autosomal coding sequence into *n* bins with length equivalent to the X-linked coding sequence, resampling estimates of the X/A ratio of $\omega_a$ across these bins, and then calculating how many resampled estimates were larger in the "Fully conserved" than the "Partly conserved" SIFT category). **I.** Correlation between X/A ratios of $\omega_a$ and Phylostratum (Spearman's $\rho$=-0.070, p=0.834).

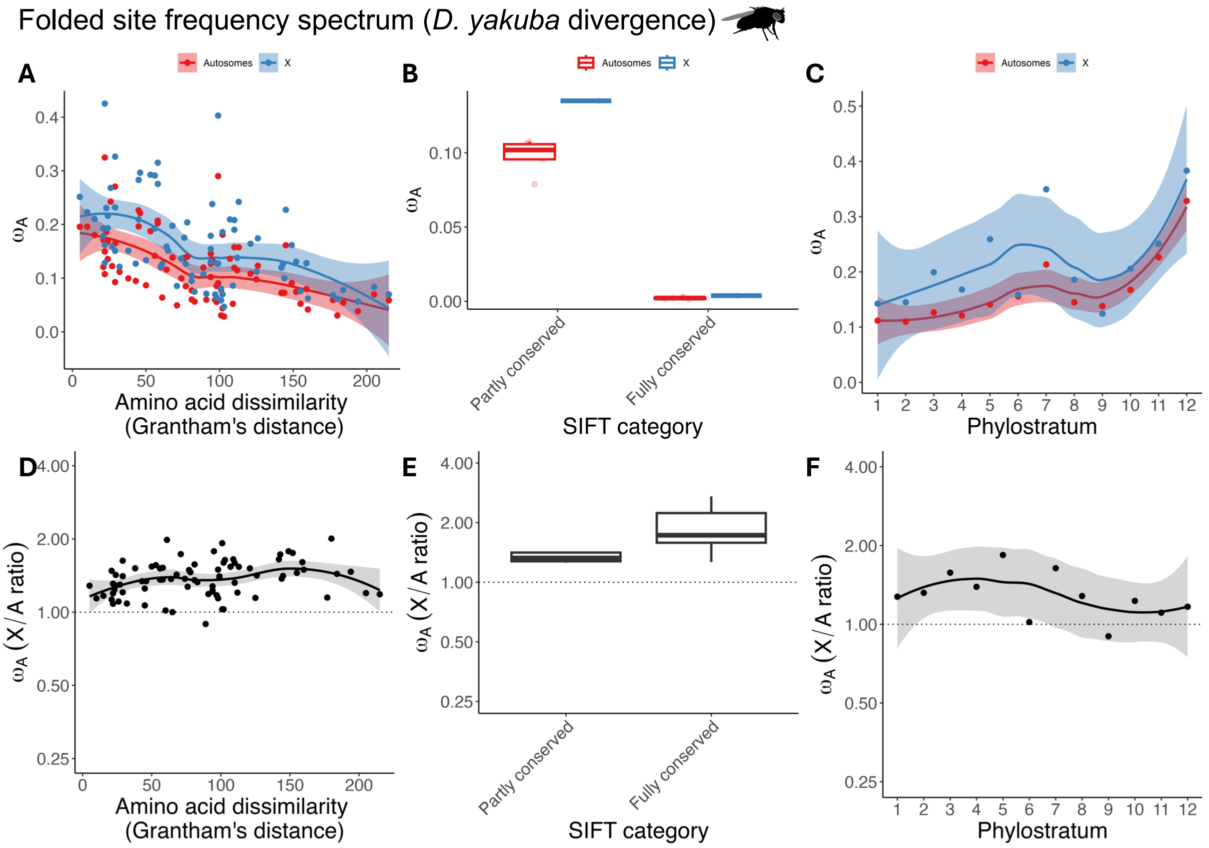


**Supplementary Figure 4. Proxies for effect size and relative rates of X-autosome adaptation in *D. melanogaster*, using the "GammaZero" model for fitting the DFE to the folded site frequency spectrum, calculated from divergence to *D. yakuba*. A.** Rates of adaptation, $\omega_a$, for nonsynonymous mutations affecting 81 pairs of amino acids, plotted against the Grantham's distance between each amino-acid pair. **B.** Same as A, but for nonsynonymous mutations categorised as having "Partly conserved" or "Fully conserved" SIFT scores. **C.** Same as A, but nonsynonymous mutations that fall in genes with different Phylostrata (smaller values indicate that genes are older). **D.** Correlation between X/A ratios of $\omega_a$ and Grantham's distance (Spearman's $\rho$=0.315, p=0.004), with X/A ratios plotted on a log2 scale. Negative values of $\omega_a$ were treated as zeroes and Infinite values were treated as $10^3$. **E.** X/A ratios of $\omega_a$ vs. SIFT score category (p=0.182, obtained by splitting the autosomal coding sequence into *n* bins with length equivalent to the X-linked coding sequence, resampling estimates of the X/A ratio of $\omega_a$ across these bins, and then calculating how many resampled estimates were larger in the "Fully conserved" than the "Partly conserved" SIFT category). **F.** Correlation between X/A ratios of $\omega_a$ and Phylostratum (Spearman's $\rho$=-0.517, p=0.089).

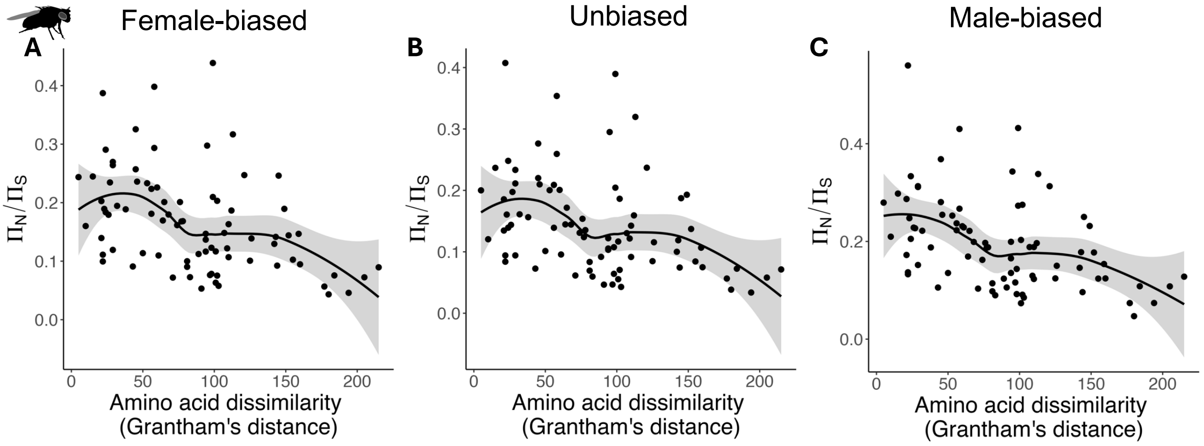


**Supplementary Figure 5. Grantham's distances and ratios of autosomal nonsynonymous to synonymous nucleotide diversity in *D. melanogaster*, for different bins of sex-biased expression.** Autosomal $\pi_N/\pi_S$ estimates, for polymorphisms affecting 81 pairs of amino acids, ranked by their Grantham's distance, fitted with a loess regression curve. Shown for female-biased genes (Spearman's $\rho$=-0.476, $p$<0.001), unbiased genes (Spearman's $\rho$=-0.435, $p$<0.001) and male-biased genes (Spearman's $\rho$=-0.463, $p$<0.001).

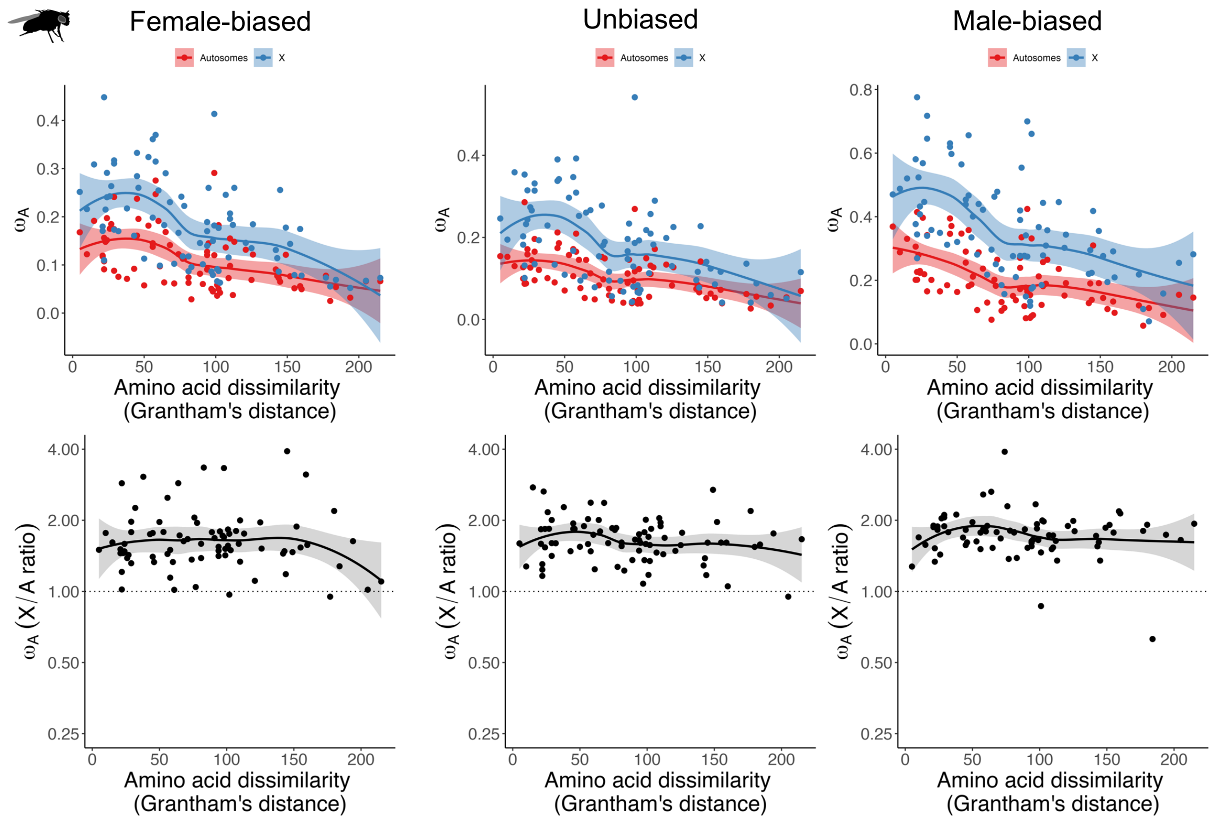


**Supplementary Figure 6. Grantham's distances and relative rates of X-autosome adaptation in *D. melanogaster*, for different bins of sex-biased expression. A-C.** Rates of adaptation, $\omega_a$, for nonsynonymous mutations affecting each pair of amino acids, plotted against the Grantham's distance between each amino-acid pair. Shown for female-biased genes, unbiased genes and male-biased genes. **D-F.** Correlation between X/A ratios of $\omega_a$ and Grantham's distance (female-biased genes: Spearman's $\rho = -0.003, p = 0.976$; unbiased genes: Spearman's $\rho = -0.123, p = 0.276$; male-biased genes: Spearman's $\rho = -0.025, p = 0.827$), with X/A ratios plotted on a log2 scale. Negative values of $\omega_a$ were treated as zeroes and Infinite values were treated as $10^3$.

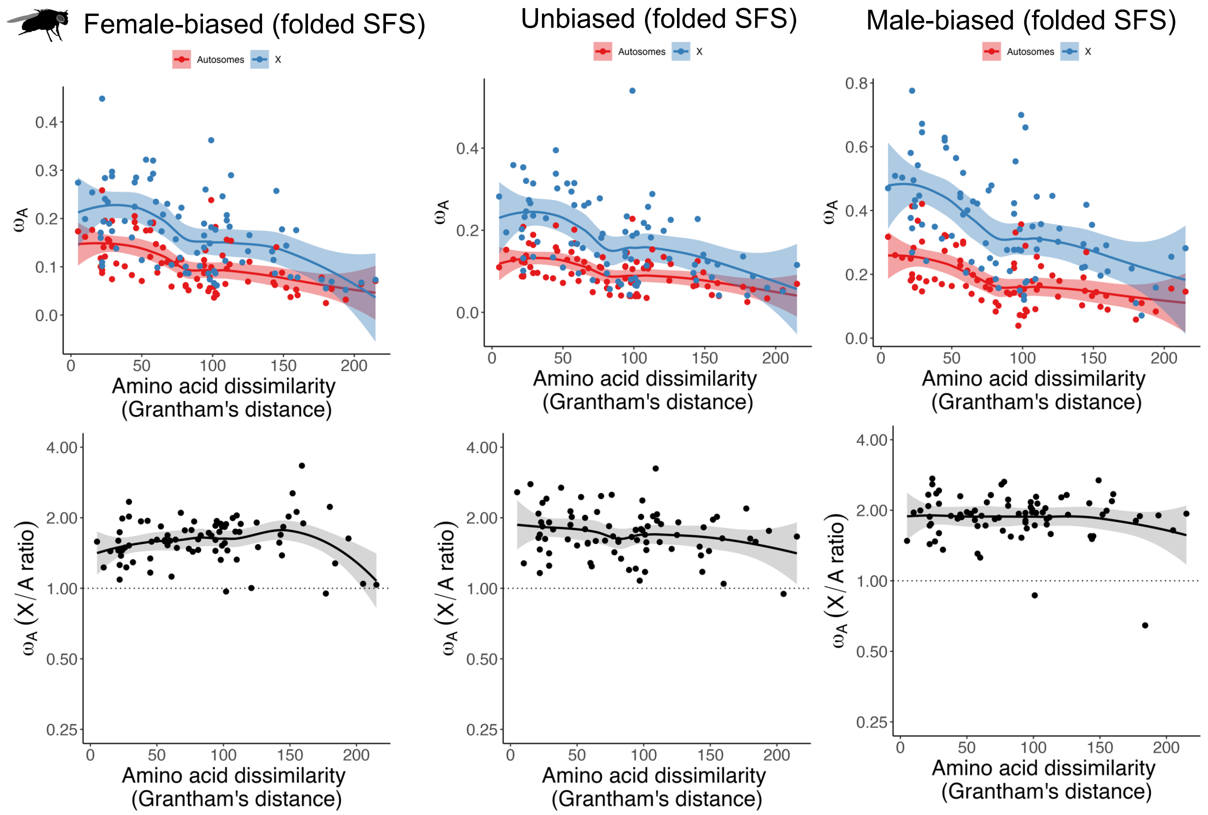


**Supplementary Figure 7. Grantham's distances and relative rates of X-autosome adaptation in *D. melanogaster*, using the "GammaZero" model for fitting the DFE to the folded site frequency spectrum, for different bins of sex-biased expression. A-C.** Rates of adaptation, $\omega_a$, for nonsynonymous mutations affecting each pair of amino acids, plotted against the Grantham's distance between each amino-acid pair. Shown for female-biased genes, unbiased genes and male-biased genes. **D-F.** Correlation between X/A ratios of $\omega_a$ and Grantham's distance (female-biased genes: Spearman's $\rho = -0.003, p = 0.976$; unbiased genes: Spearman's $\rho = -0.123, p = 0.276$; male-biased genes: Spearman's $\rho = -0.025, p = 0.827$), with X/A ratios plotted on a log2 scale. Negative values of $\omega_a$ were treated as zeroes and Infinite values were treated as $10^3$.

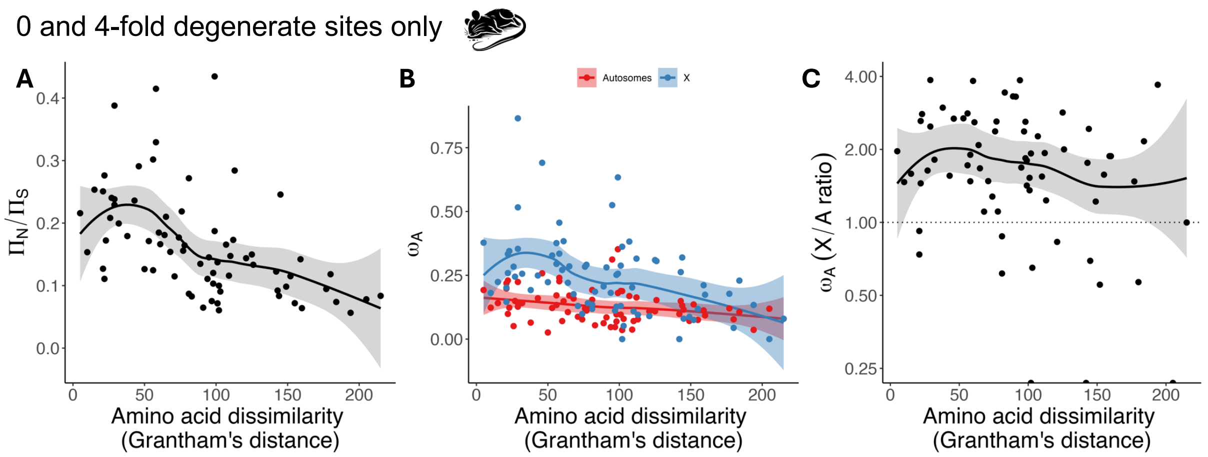


**Supplementary Figure 8. Grantham's distances, ratios of autosomal nonsynonymous to synonymous nucleotide diversity, and relative rates of X-autosome adaptation in *M. musculus*. A.** Autosomal $\pi_0/\pi_4$ estimates, for polymorphisms affecting each pair of amino acid, ranked by Grantham's distance, fitted with a loess regression curve. A significant negative rank correlation was observed (Spearman's $\rho$=-0.596, $p<0.001$). **B.** Rates of adaptation, $\omega_a$, for nonsynonymous mutations affecting each pair of amino acids, plotted against the Grantham's distance between each amino-acid pair. **C.** Correlation between X/A ratios of $\omega_a$ and Grantham's distance (Spearman's $\rho = -0.277, p = 0.018$), with X/A ratios plotted on a log2 scale. Negative values of $\omega_a$ were treated as zeroes and Infinite values were treated as $10^3$.

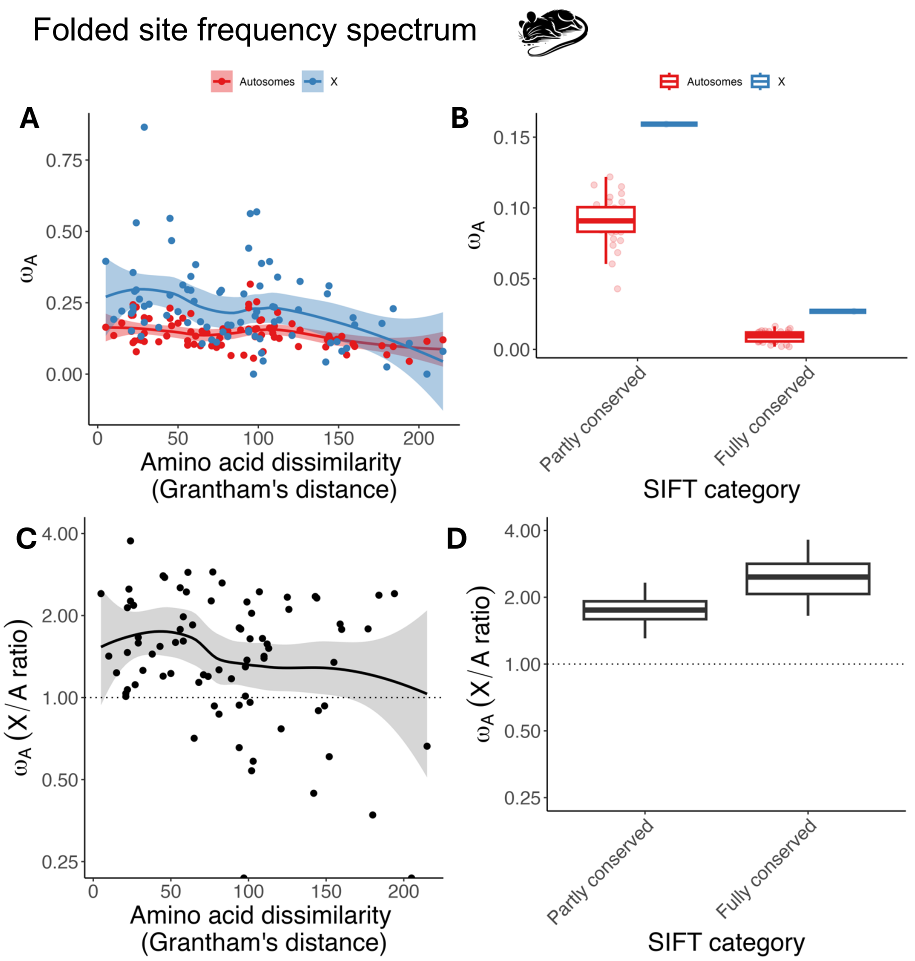


**Supplementary Figure 9. Proxies for effect size and relative rates of X-autosome adaptation in *M. musculus*, using the "GammaZero" model for fitting the DFE to the folded site frequency spectrum. A.** Rates of adaptation, $\omega_a$, for nonsynonymous mutations affecting 81 pairs of amino acids, plotted against the Grantham's distance between each amino-acid pair. **B.** Same as A, but for nonsynonymous mutations categorised as having "Partly conserved" or "Fully conserved" SIFT scores. **C.** Correlation between X/A ratios of $\omega_a$ and Grantham's distance (Spearman's ρ=-0.205, p=0.066), with X/A ratios plotted on a log2 scale. Negative values of $\omega_a$ were treated as zeroes and Infinite values were treated as $10^3$. **D.** X/A ratios of $\omega_a$ vs. SIFT score category (p=0.081, obtained by splitting the autosomal coding sequence into *n* bins with length equivalent to the X-linked coding sequence, resampling estimates of the X/A ratio of $\omega_a$ across these bins, and then calculating how many resampled estimates were larger in the "Fully conserved" than the "Partly conserved" SIFT category).

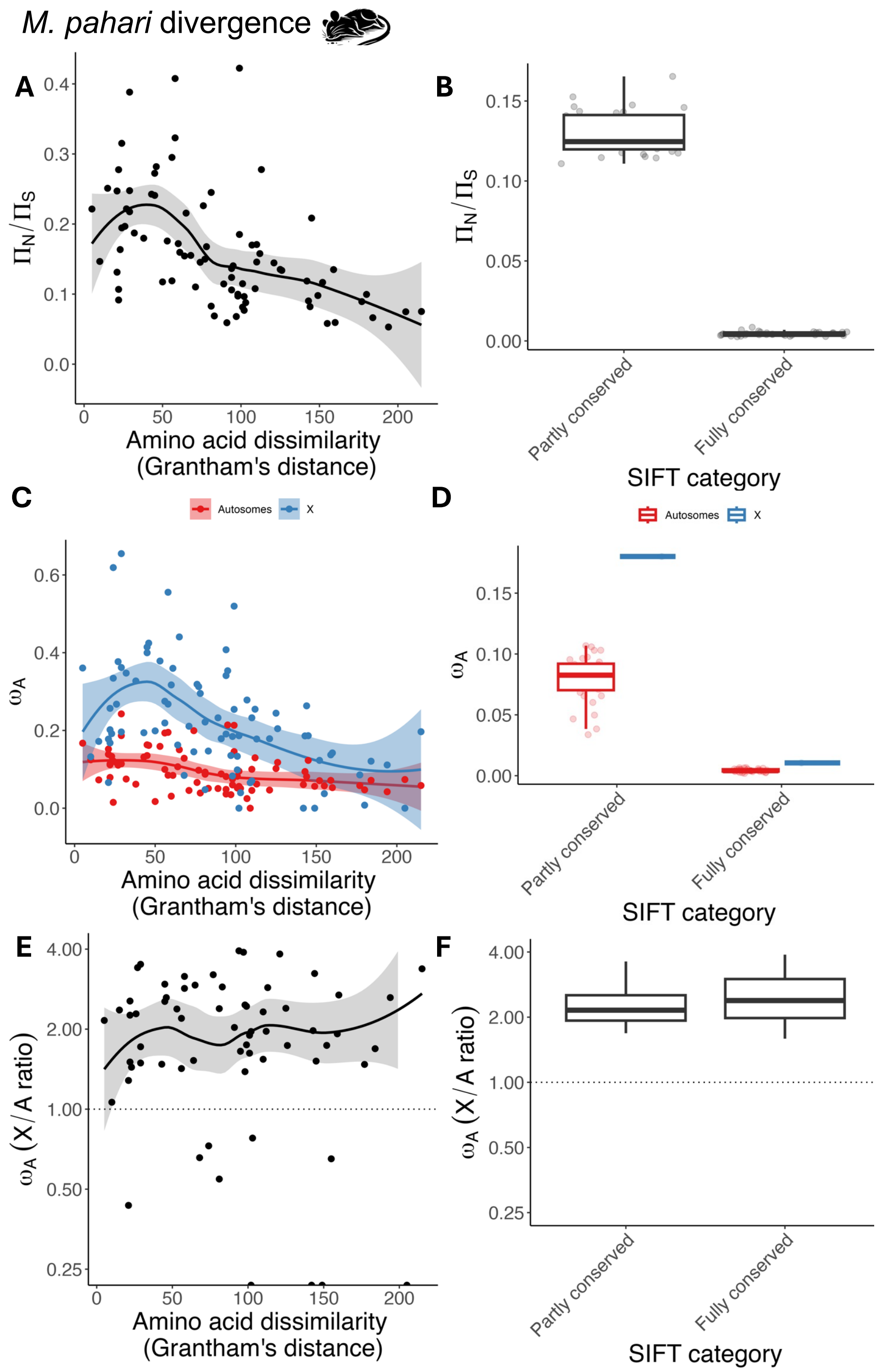


**Supplementary Figure 10. Proxies for effect size, ratios of autosomal nonsynonymous to synonymous nucleotide diversity, and relative rates of X-autosome adaptation in *M. musculus*, calculated from divergence to *M. pahari*. A.** Autosomal $\pi_N/\pi_S$ estimates, for polymorphisms affecting 81 pairs of amino acids, ranked by their Grantham's distance, fitted with a loess regression curve. A significant negative rank correlation was observed (Spearman's $\rho$=-0.591, $p$<0.001). **B.** Same as A, but autosomal $\pi_N/\pi_S$ is compared between "Partly conserved" and "Fully conserved" SIFT score categories ($p$<0.001, based on splitting the autosomal coding sequence into n bins with length equivalent to the X-linked coding sequence, and resampling autosomal estimates across these bins). **C.** Rates of adaptation, $\omega_a$, for nonsynonymous mutations affecting 81 pairs of amino acids, plotted against the Grantham's distance between each amino-acid pair. **D.** Same as C, but for nonsynonymous mutations categorised as having "Partly conserved" or "Fully conserved" SIFT scores. **E.** Correlation between X/A ratios of $\omega_a$ and Grantham's distance (Spearman's $\rho$=-0.161, $p<0.001$), with X/A ratios plotted on a log2 scale. Negative values of $\omega_a$ were treated as zeroes and Infinite values were treated as $10^3$. **F.** X/A ratios of $\omega_a$ vs. SIFT score category ($p$=0.401, obtained by splitting the autosomal coding sequence into *n* bins with length equivalent to the X-linked coding sequence, resampling estimates of the X/A ratio of $\omega_a$ across these bins, and then calculating how many resampled estimates were larger in the "Fully conserved" than the "Partly conserved" SIFT category).

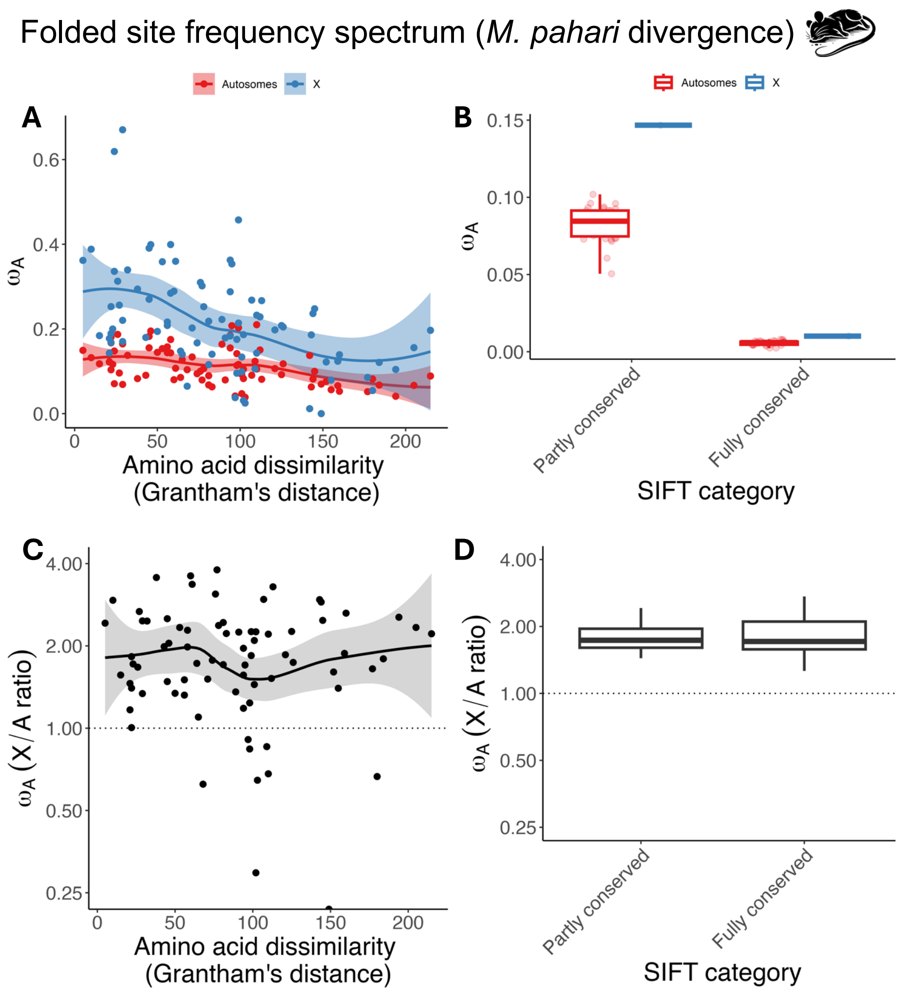


**Supplementary Figure 11. Proxies for effect size and relative rates of X-autosome adaptation in *M. musculus*, using the "GammaZero" model for fitting the DFE to the folded site frequency spectrum, calculated from divergence to *M. pahari*. A.** Rates of adaptation, $\omega_a$, for nonsynonymous mutations affecting 81 pairs of amino acids, plotted against the Grantham's distance between each amino-acid pair. **B.** Same as A, but for nonsynonymous mutations categorised as having "Partly conserved" or "Fully conserved" SIFT scores. **C.** Correlation between X/A ratios of $\omega_a$ and Grantham's distance (Spearman's $\rho$=-0.107, p=0.340), with X/A ratios plotted on a log2 scale. Negative values of $\omega_a$ were treated as zeroes and Infinite values were treated as $10^3$. **D.** X/A ratios of $\omega_a$ vs. SIFT score category (p=0.508, obtained by splitting the autosomal coding sequence into *n* bins with length equivalent to the X-linked coding sequence, resampling estimates of the X/A ratio of $\omega_a$ across these bins, and then calculating how many resampled estimates were larger in the "Fully conserved" than the "Partly conserved" SIFT category).

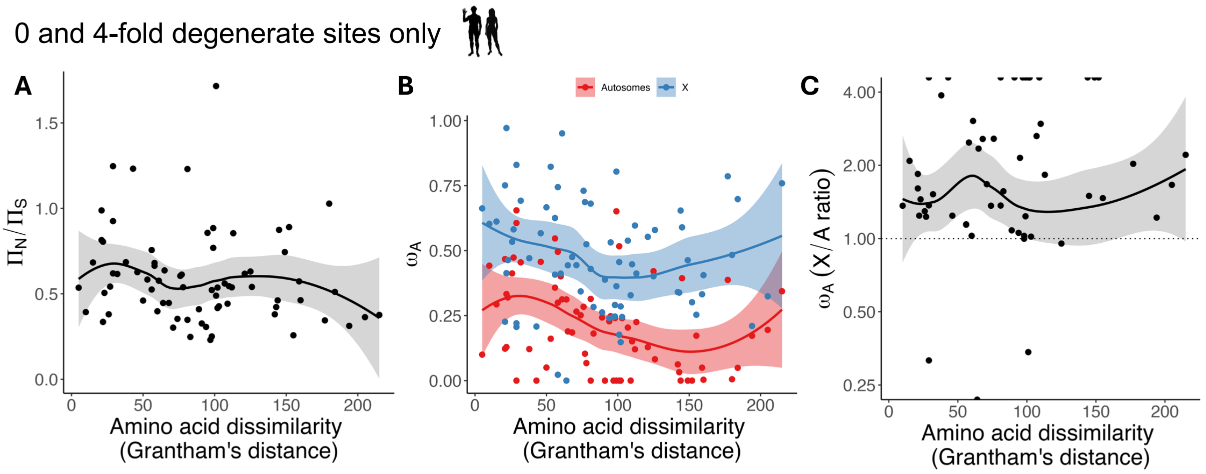


**Supplementary Figure 12. Grantham's distances, ratios of autosomal nonsynonymous to synonymous nucleotide diversity, and relative rates of X-autosome adaptation in *H. sapiens*. A.** Autosomal $\pi_0/\pi_4$ estimates, for polymorphisms affecting each pair of amino acid, ranked by Grantham's distance, fitted with a loess regression curve. A non-significant negative rank correlation was observed (Spearman's ρ=-0.199, p=0.074). **B.** Rates of adaptation, $\omega_a$, for nonsynonymous mutations affecting each pair of amino acids, plotted against the Grantham's distance between each amino-acid pair. **C.** Correlation between X/A ratios of $\omega_a$ and Grantham's distance (Spearman's $\rho = 0.252, p = 0.031$), with X/A ratios plotted on a log2 scale. Negative values of $\omega_a$ were treated as zeroes and Infinite values were treated as $10^3$.

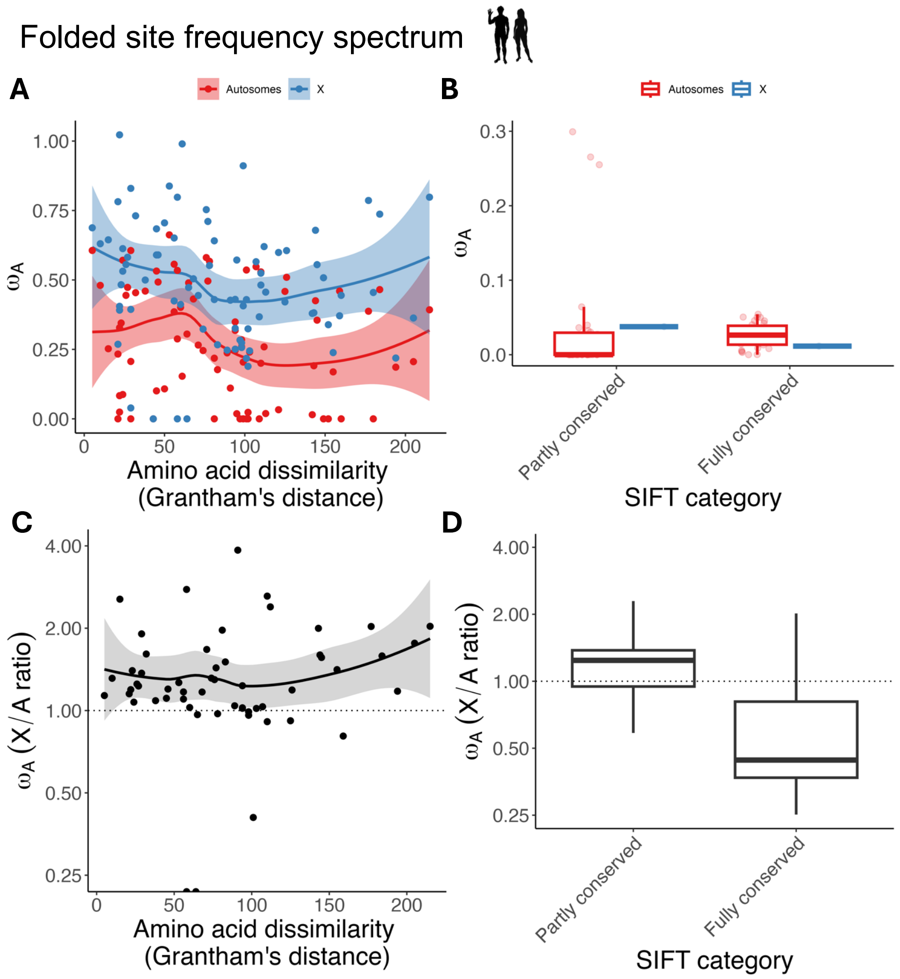


**Supplementary Figure 13. Proxies for effect size and relative rates of X-autosome adaptation in *H. sapiens*, using the "GammaZero" model for fitting the DFE to the folded site frequency spectrum. A.** Rates of adaptation, $\omega_a$, for nonsynonymous mutations affecting 81 pairs of amino acids, plotted against the Grantham's distance between each amino-acid pair. **B.** Same as A, but for nonsynonymous mutations categorised as having "Partly conserved" or "Fully conserved" SIFT scores. **C.** Correlation between X/A ratios of $\omega_a$ and Grantham's distance (Spearman's $\rho$=0.147, p=0.193), with X/A ratios plotted on a log2 scale. Negative values of $\omega_a$ were treated as zeroes and Infinite values were treated as $10^3$. **D.** X/A ratios of $\omega_a$ vs. SIFT score category (p=0.800, obtained by splitting the autosomal coding sequence into *n* bins with length equivalent to the X-linked coding sequence, resampling estimates of the X/A ratio of $\omega_a$ across these bins, and then calculating how many resampled estimates were larger in the "Fully conserved" than the "Partly conserved" SIFT category).

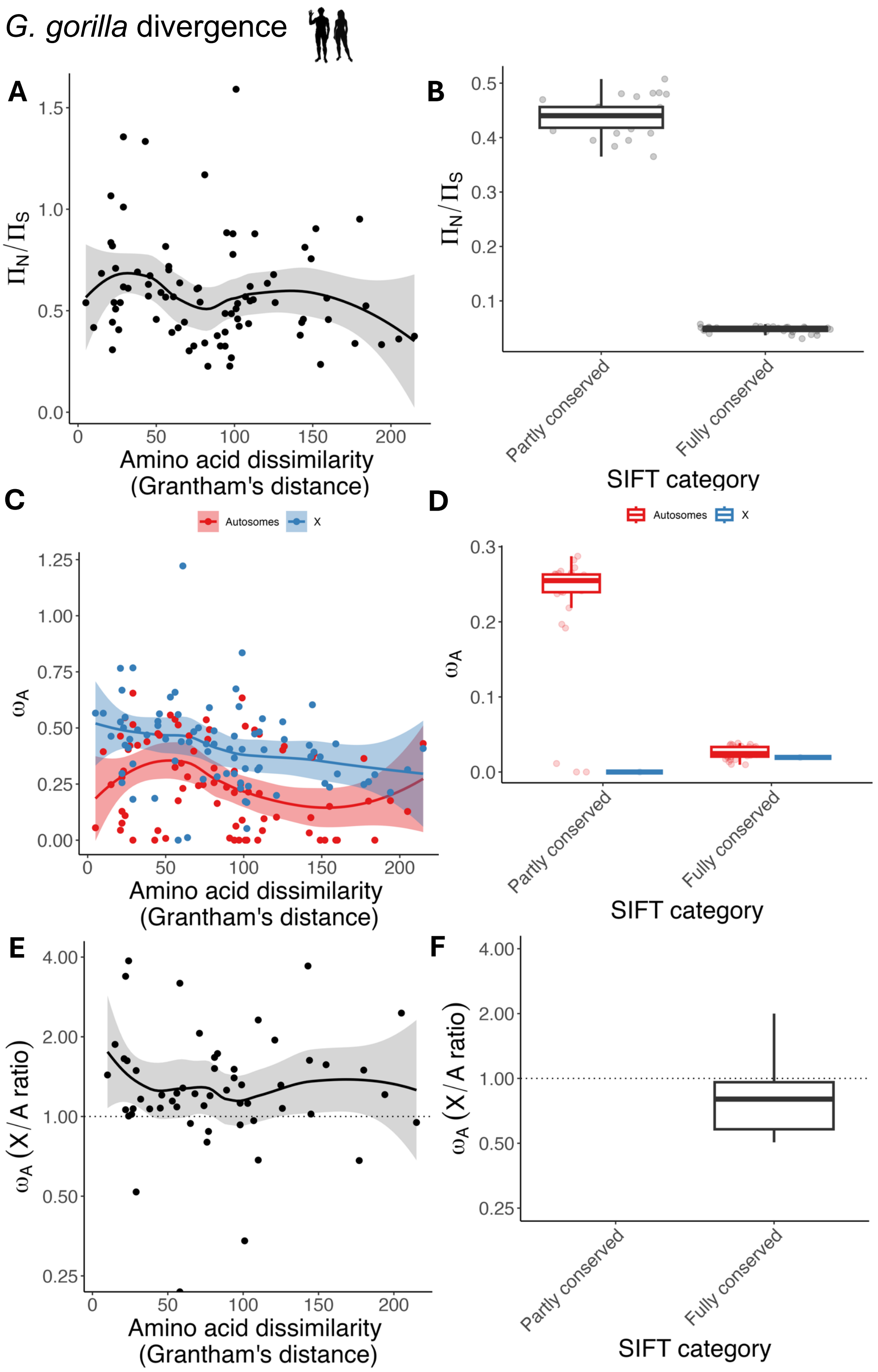


**Supplementary Figure 14. Proxies for effect size, ratios of autosomal nonsynonymous to synonymous nucleotide diversity, and relative rates of X-autosome adaptation in *H. sapiens*, calculated from divergence to *G. gorilla*. A.** Autosomal $\pi_N/\pi_S$ estimates, for polymorphisms affecting 81 pairs of amino acids, ranked by their Grantham's distance, fitted with a loess regression curve. A non-significant negative rank correlation was observed (Spearman's $\rho$=-0.195, p=0.082). **B.** Same as A, but autosomal $\pi_N/\pi_S$ is compared between "Partly conserved" and "Fully conserved" SIFT score categories (p<0.001, based on splitting the autosomal coding sequence into n bins with length equivalent to the X-linked coding sequence, and resampling autosomal estimates across these bins). **C.** Rates of adaptation, $\omega_a$, for nonsynonymous mutations affecting 81 pairs of amino acids, plotted against the Grantham's distance between each amino-acid pair. **D.** Same as C, but for nonsynonymous mutations categorised as having "Partly conserved" or "Fully conserved" SIFT scores. **E.** Correlation between X/A ratios of $\omega_a$ and Grantham's distance (Spearman's $\rho=0.139$, $p=0.215$), with X/A ratios plotted on a log2 scale. Negative values of $\omega_a$ were treated as zeroes and Infinite values were treated as $10^3$. **F.** X/A ratios of $\omega_a$ vs. SIFT score category ($p<0.001$, obtained by splitting the autosomal coding sequence into *n* bins with length equivalent to the X-linked coding sequence, resampling estimates of the X/A ratio of $\omega_a$ across these bins, and then calculating how many resampled estimates were larger in the "Fully conserved" than the "Partly conserved" SIFT category). Note that the estimated X/A ratios for the "Partly conserved" category are ~0, because almost no adaptation was detected for the X chromosome in this subset of nonsynonymous sites (see panel D).

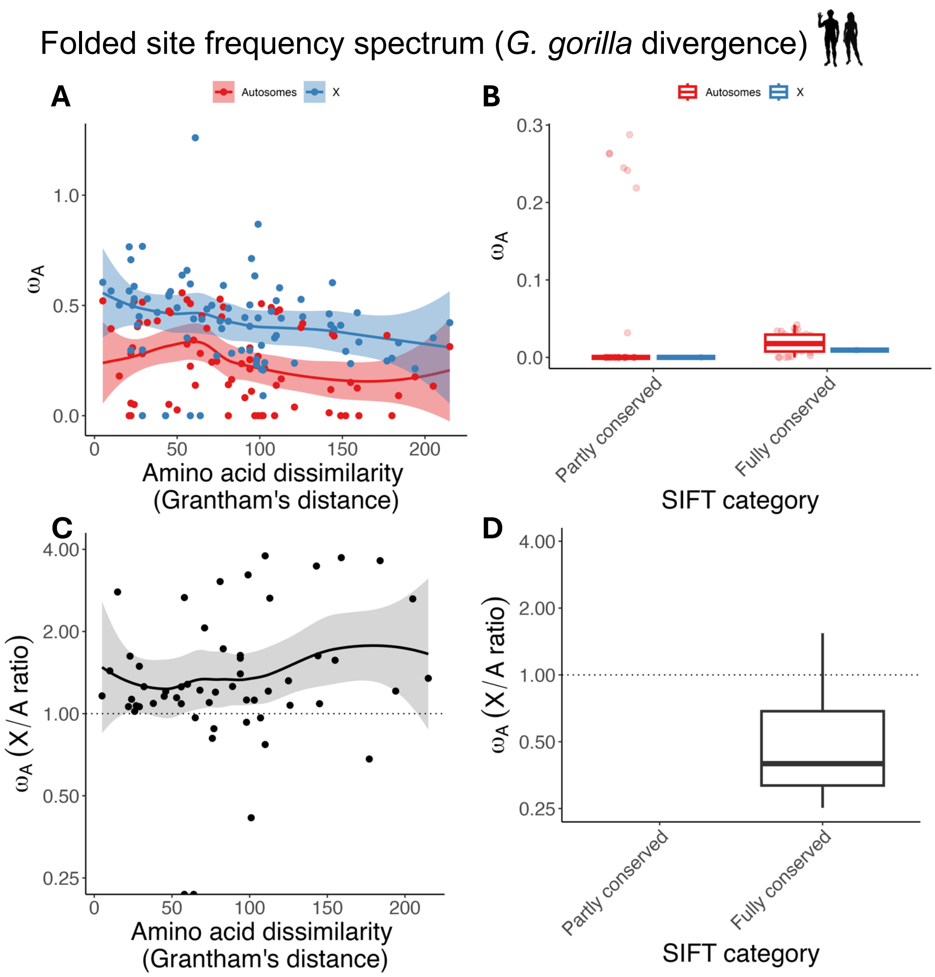


**Supplementary Figure 15. Proxies for effect size and relative rates of X-autosome adaptation in *H. sapiens*, using the "GammaZero" model for fitting the DFE to the folded site frequency spectrum, calculated from divergence to *G. gorilla*. A.** Rates of adaptation, $\omega_a$, for nonsynonymous mutations affecting 81 pairs of amino acids, plotted against the Grantham's distance between each amino-acid pair. **B.** Same as A, but for nonsynonymous mutations categorised as having "Partly conserved" or "Fully conserved" SIFT scores. **C.** Correlation between X/A ratios of $\omega_a$ and Grantham's distance (Spearman's $\rho$=0.148, p=0.195), with X/A ratios plotted on a log2 scale. Negative values of $\omega_a$ were treated as zeroes and Infinite values were treated as $10^3$. **D.** X/A ratios of $\omega_a$ vs. SIFT score category (p<0.001, obtained by splitting the autosomal coding sequence into *n* bins with length equivalent to the X-linked coding sequence, resampling estimates of the X/A ratio of $\omega_a$ across these bins, and then calculating how many resampled estimates were larger in the "Fully conserved" than the "Partly conserved" SIFT category). Note that the estimated X/A ratios for the "Partly conserved" category are ~0, because almost no adaptation was detected for the X chromosome in this subset of nonsynonymous sites (see panel B).

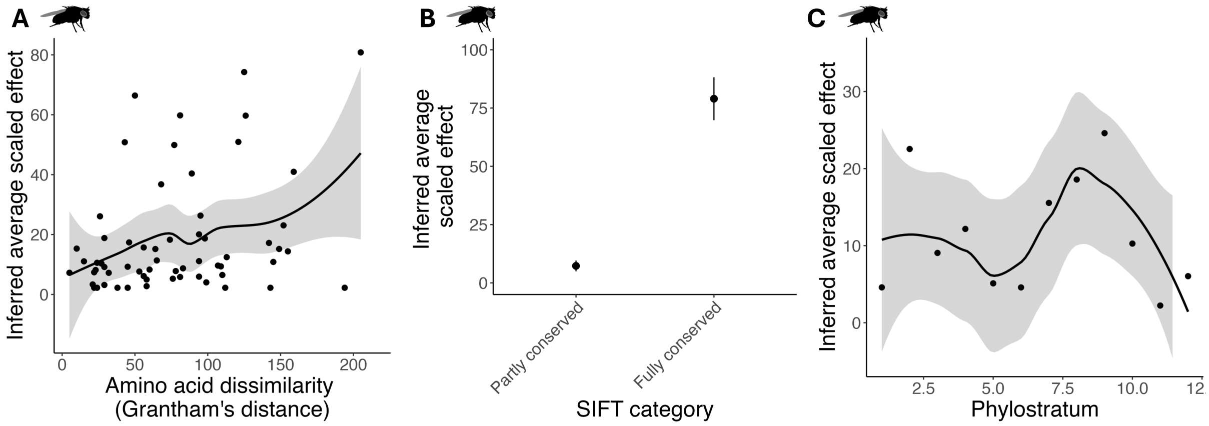


**Supplementary Figure 16. The average scaled phenotypic effect size (*x*) inferred from the proportion of mutations that are positively selected (Fig. 3D-F) on autosomes in *D. melanogaster*.**

Expressions that relate values of average $x$ ($\bar{x}$) to the proportion of mutations that are positively selected were derived as follows. Assuming phenotypic dominance has a constant value $v$ and mutational effects conform to the standard isotropic version of Fisher's geometric model (Fisher 1930; Orr 1998) with relatively high dimensionality (i.e., $n > \sim 10$, where $n$ is the number of trait dimensions), the probability that a random autosomal mutation with effect size $x$ is subject to positive selection is:

$$\Pr(\text{pos.}|A, x) \approx \frac{1}{2}\left(1 - \text{erf}\left(\frac{x(1+v)}{\sqrt{2}}\right)\right)$$

following McDonough et al. (2024). Assuming that the distribution of scaled effect sizes is exponential with a mean of $\bar{x}$, then the total probability that a random autosomal mutation is under positive selection is:

$$\Pr(\text{pos.}|A) \approx \int_0^\infty \Pr(\text{pos.}|A, x)\, f(x)dx \approx \int_0^\infty \frac{1}{2}\left(1 - \text{erf}\left(\frac{x(1+v)}{\sqrt{2}}\right)\right)\frac{1}{\bar{x}}\exp\left(-\frac{x}{\bar{x}}\right)dx$$

$$= \frac{1}{2}\left(1 - \frac{1}{\bar{x}}\int_0^\infty \text{erf}\left(\frac{x(1+v)}{\sqrt{2}}\right)\exp\left(-\frac{x}{\bar{x}}\right)dx\right)$$

$$= \frac{1 - \exp\left(\frac{1}{2}(1+v)^{-2}\bar{x}^{-2}\right)\left(1 - \text{erf}\left(\frac{(1+v)^{-1}\bar{x}^{-1}}{\sqrt{2}}\right)\right)}{2}$$

This matches eq. (15) of McDonough and Connallon (2023) for the special case of $v = 0$. The figure above shows results for autosomes. The proportion of random X-linked mutations that are subject to positive selection can also be related to $\bar{x}$ using the following expression:

$$\Pr(\text{pos.}|X) \approx \frac{1 - \exp\left(\frac{1}{2}(1+\psi)^{-2}\bar{x}^{-2}\right)\left(1 - \text{erf}\left(\frac{(1+\psi)^{-1}\bar{x}^{-1}}{\sqrt{2}}\right)\right)}{2}$$

where $\psi = \frac{2v(1-v)}{3-2v}$.
